# A natural single-guide RNA repurposes Cas9 to autoregulate CRISPR-Cas expression

**DOI:** 10.1101/2020.05.21.102756

**Authors:** Rachael E. Workman, Teja Pammi, Binh T. K. Nguyen, Leonardo W. Graeff, Erika Smith, Suzanne M. Sebald, Marie J. Stoltzfus, Joshua W. Modell

**Affiliations:** Department of Molecular Biology & Genetics, Johns Hopkins University School of Medicine, Baltimore, MD 21205, USA; Department of Biological Chemistry, Johns Hopkins University School of Medicine, Baltimore, MD 21205, USA

**Keywords:** CRISPR-Cas, regulation, Cas9, tracrRNA, natural single guide, bacteriophage

## Abstract

CRISPR-Cas systems provide their prokaryotic hosts with acquired immunity against viruses and other foreign genetic elements, but how these systems are regulated to prevent auto-immunity is poorly understood. In type II CRISPR-Cas systems, a transactivating CRISPR RNA (tracrRNA) scaffold functions together with a CRISPR RNA (crRNA) guide to program Cas9 for the recognition and cleavage of foreign DNA targets. Here, we show that a long-form tracrRNA performs an unexpected second function by folding into a natural single guide that directs Cas9 to transcriptionally repress its own promoter. Further, we demonstrate that P_cas9_ serves as a critical regulatory node; de-repression causes a dramatic induction of Cas genes, crRNAs and tracrRNAs resulting in a 3,000-fold increase in immunization rates against unrecognized viruses. Heightened immunity comes at the cost of increased auto-immune toxicity, demonstrating the critical importance of the controller. Using a bioinformatic analysis, we provide evidence that tracrRNA-mediated autoregulation is widespread in type II CRISPR-Cas systems. Collectively, we unveil a new paradigm for the intrinsic regulation of CRISPR-Cas systems by natural single guides, which may facilitate the frequent horizontal transfer of these systems into new hosts that have not yet evolved their own regulatory strategies.

## INTRODUCTION

All immune systems must distinguish “self” from “non-self” in order to provide potent activity against diverse pathogen-associated antigens while avoiding autoimmunity against the more abundant molecular catalog of the host. To help ensure specificity against foreign threats, the vertebrate adaptive immune system is a tightly regulated cellular network that undergoes spatiotemporal activation and expansion in response to novel or remembered antigens^1^. CRISPR-Cas systems provide bacteria and archaea with immunological memories of foreign nucleic acids, thereby protecting their prokaryotic hosts from viruses^2^, plasmids^3^, and other mobile genetic elements^4,5^. How CRISPR-Cas immune systems are regulated within their single-celled hosts is not well understood.

Immunological memories are encoded within CRISPR loci as short, roughly 30 bp DNA “spacers” which are derived from segments of foreign nucleic acids^2^. These spacers are located within CRISPR arrays between similarly-sized repeating sequences or “repeats”^6^. CRISPR immunity occurs in three stages. During “adaptation,” spacers are acquired from foreign agents and incorporated into the CRISPR array at the end proximal to a conserved “leader” sequence, causing the duplication of the leader-proximal repeat^2^. Under laboratory conditions, adaptation is a rare event, occurring in roughly 1E-5 to 1E-7 cells^7,8^. During “biogenesis,” the CRISPR array is transcribed as one long precursor CRISPR RNA (pre-crRNA), which is cleaved within repeats to produce individual CRISPR RNAs (crRNAs), each containing a single spacer^9–11^. Finally, during “interference”, crRNAs direct effector Cas proteins to matching foreign targets^12–14^, where they perform distinct catalytic activities depending on the CRISPR-Cas family and sub-type^15^. The type II-A CRISPR-Cas system from *Streptococcus pyogenes* encodes four Cas proteins, expressed as a single operon, all of which are required for adaptation (Fig. 1A)^7^. However, Cas9 alone performs interference by introducing double-strand breaks into crRNA-specified DNA targets or “protospacers”^16,17^. To prevent the CRISPR system from cleaving its own spacers, Cas9 targeting requires a 5’-NGG-3’ protospacer-adjacent motif (PAM)^18,19^ which is absent in the spacer-adjacent repeats within the CRISPR array. Nonetheless, at high concentrations, Cas9 can cleave off-target genomic loci suggesting that additional PAM-independent regulatory mechanisms may be required to avoid auto-immunity.

**Figure 1.**
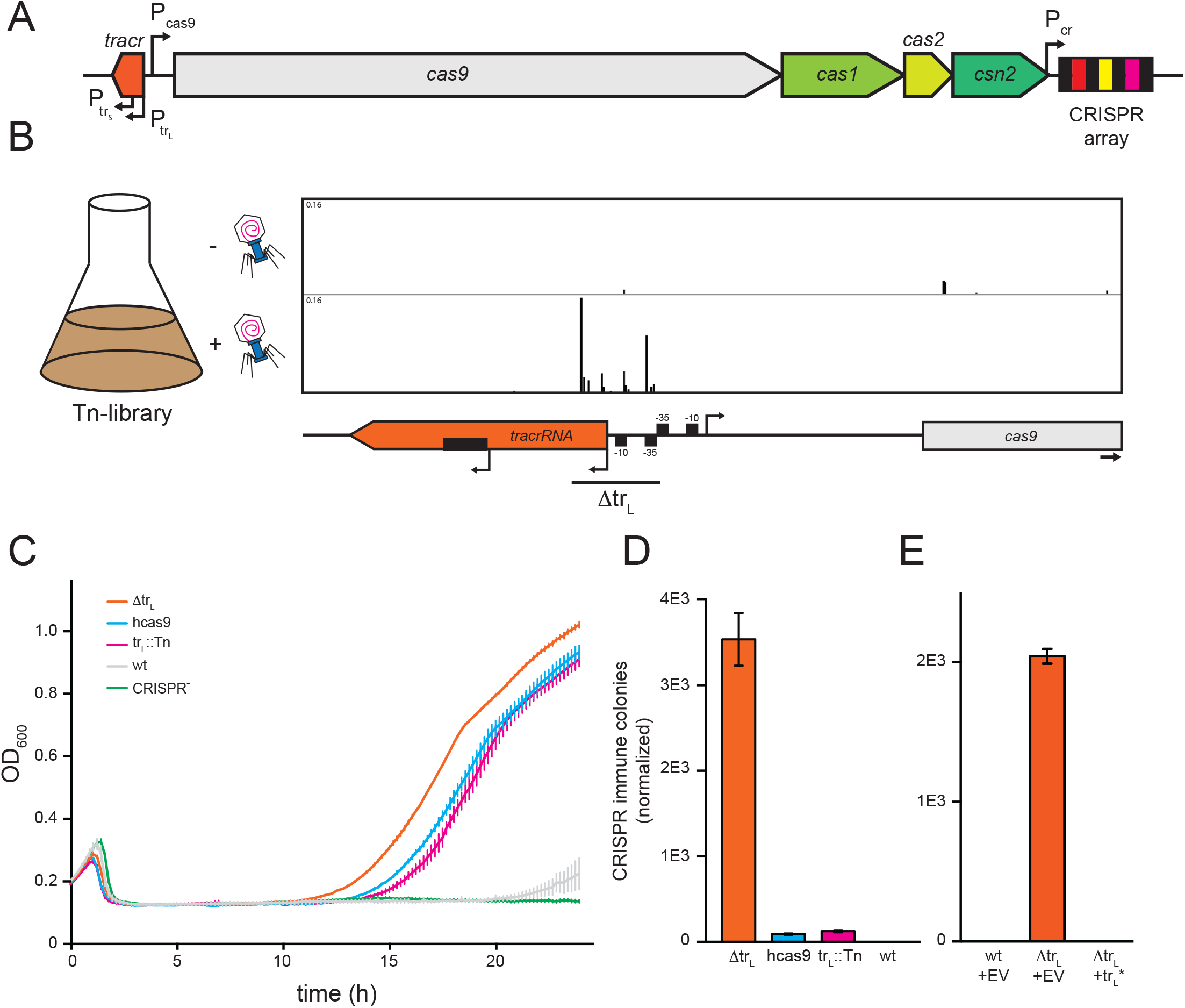
Tn-seq reveals a tracrRNA mutant with enhanced CRISPR immunity. **A)** Schematic of the *S. pyogenes* type II CRISPR-Cas system. Arrows indicate promoters. CRISPR array repeats (black boxes) and spacers (colored boxes) are shown. **B)** Tn-seq reads are overlaid on the corresponding tracrRNA and P_cas9_ DNA regions. NGS reads at each site are shown as the fraction of total reads. Data is representative of biological replicates. Top panel, - phage; bottom panel, + phage. Horizontal bar indicates the region deleted in Δtr_L_. **C)** Cells harboring naïve CRISPR-Cas systems with the indicated mutations were infected with ϕNM4γ4 at MOI = 10 and cell densities (OD_600_) were measured every 10 minutes in a 96-well plate reader. hcas9, hyper-Cas9; tr_L_::Tn, transposon insertion in P_tracr_; CRISPR^−^, no CRISPR system. Error bars represent SEM. **D-E)** Naïve CRISPR-Cas mutants were infected with ϕNM4γ4 at MOI = 25 in top agar, and surviving colonies with expanded CRISPR arrays were quantified. Bar graphs here and throughout show mean +/-SEM. **E)** An empty vector (EV) or a plasmid expressing long-form tracrRNA (tr_L_*) was introduced to the indicated strains from (D).

In addition to the pre-crRNA, the *S. pyogenes* CRISPR-Cas system harbors a second non-coding RNA, the tracrRNA, which functions during all three stages of CRISPR immunity^7,11^. While its role during adaptation is unclear, during biogenesis, the tracrRNA base-pairs with repeat-derived sequences within the pre-crRNA, and the tracrRNA:pre-crRNA duplex is then cleaved by the host factor RNaseIII within each repeat, producing individual crRNAs and a processed tracrRNA (tr_P_)^11^ (Fig. S1). During interference, the mature targeting complex (Cas9:tr_P_:crRNA) facilitates target recognition and cleavage. Many CRISPR editing technologies employ an artificial single-guide RNA (sgRNA) in which tr_P_ and crRNA are fused with a 5’-GAAA-3’ tetraloop^13^. In *S. pyogenes*, the tracrRNA is transcribed from two promoters producing a long (tr_L_) and short (tr_S_) form, both of which contain the RNaseIII processing site (Fig. 1B, S1)^11^. Because tr_S_ alone can mediate crRNA biogenesis and interference *in vivo* and *in vitro*, it remains unclear what role tr_L_ plays and why two tracrRNA forms are produced.

How are CRISPR systems regulated within their bacterial hosts to prevent autoimmunity and enhance targeting of foreign agents? Several studies have demonstrated transcriptional and post-transcriptional CRISPR-Cas regulation in response to extracellular and intracellular cues^20^, including phage infection^21–25^, quorum sensing^26,27^, membrane stress^28–31^, metabolic status^21,32,33^ and surface association^34^. In many cases where the regulatory mechanisms are known, host-encoded transcription factors interact with CRISPR-Cas promoters^32,33,35,36^, although several studies have shown that archaeal type I-A systems can be intrinsically controlled by dedicated Cas-encoded transcription factors^25,37–40^. Given that CRISPR systems are frequently horizontally transferred in the wild^41–43^, these intrinsic controllers could help prevent autoimmune toxicity^44–47^ in a new host that has not had time to evolve its own regulatory strategy. However, few CRISPR loci - and no type II loci - encode dedicated transcription factors, and it is unclear whether alternative mechanisms of intrinsic regulation exist.

Here, we identify a novel mechanism of intrinsic control, whereby a long-form tracrRNA (tr_L_) enables Cas9 to autoregulate type II CRISPR-Cas systems. Using a transposon screen, we identify tr_L_ as a potent inhibitor of CRISPR immunity. We find that tr_L_ folds into a natural single guide RNA by providing the structural elements normally contributed by the crRNA. In place of a spacer, tr_L_ contains a truncated 11 nt targeting sequence that directs Cas9 to transcriptionally silence its own promoter. We show that P_cas9_ is a critical, system-wide regulatory node that controls the RNA levels of all CRISPR-Cas components. De-repression is a double-edged sword; a 3,000-fold increase in phage defense is accompanied by the deleterious effects of increased auto-immunity. Finally, we conduct a bioinformatic analysis and observe conservation of tracrRNA regulation in many type II-A CRISPR loci. Our work highlights an unexpected function for tracrRNAs and a novel form of intrinsic control within the CRISPR-Cas module, which may facilitate the frequent horizontal transfer of CRISPR systems between hosts in the wild.

## RESULTS

### Tn-seq reveals a tracrRNA mutant with enhanced CRISPR immunity

We initially sought to identify bacterial genes involved in CRISPR-Cas immunity through an unbiased genetic screen. In our model system, *Staphylococcus aureus* cells that do not encode a native CRISPR-Cas system harbor the *S. pyogenes* type II CRISPR-Cas system on a medium or low-copy plasmid^7^. The *S. aureus* host provides genetic tractability, an assortment of well-characterized phages, and a compatible cellular environment for the *S. pyogenes* CRISPR-Cas system. Furthermore, given that CRISPR-Cas systems are frequently horizontally transferred between bacterial hosts in the wild, our heterologous expression system serves as a model for a recent transfer event.

To create an unbiased library of genomic mutants, we introduced the transposon Tn917^48^ into *S. aureus* cells carrying the *S. pyogenes* CRISPR-Cas system on a medium-copy plasmid. The CRISPR system in these cells contained a minimal “naïve” CRISPR array harboring a single CRISPR repeat and no spacers. Therefore, protection against phages required all three stages of CRISPR immunity, beginning with the acquisition of a new spacer. We next infected this library with the bacteriophage ϕNM4γ4^7^ at a multiplicity of infection (MOI) of 1 for 24 hours and identified mutants that were enriched or depleted relative to an un-infected control by deep sequencing Tn insertion sites (Tn-seq^49^). Mutations that are enriched during the infection represent enhancers of CRISPR immunity, while those that are depleted represent CRISPR repressors.

To our surprise, the transposon insertions that most strongly enhanced CRISPR immunity were within the CRISPR locus itself, spanning the 5’ end and promoter of tr_L_ (Fig. 1B, S2A, Table S1). To validate the mutant phenotype, we isolated a single phage-resistant colony and confirmed the presence of a transposon insertion within the promoter of tr_L_ (tr_L_::Tn) and a newly acquired spacer matching ϕNM4γ4. We next replaced the ϕNM4γ4-targeting CRISPR array with a single repeat in order to test the effects of tr_L_::Tn on a ‘naïve’ CRISPR system with no immunological history. In liquid and semi-solid phage immunity assays, we found that tr_L_::Tn enhances CRISPR immunity by roughly 100-fold, similar to a previously identified mutation in *cas9* (*hcas9*)^47^ that boosts spacer acquisition rates (Fig. 1C-D, S2B). To determine whether tr_L_::Tn is a gain or loss-of-function mutation, we deleted a 52 bp region spanning the CRISPR immunity-enhancing transposon insertion sites found by Tn-seq, which includes the promoter and 19 nucleotides at the 5’ end of tr_L_, generating Δtr_L_ (Fig. 1B). Strikingly, Δtr_L_ significantly enhances immunity to ϕNM4γ4 by roughly 30-fold relative to *hcas9* and tr_L_::Tn and over 3,000-fold relative to the wild-type system (Fig. 1C-D, S2B). To test whether expression of tr_L_ in *trans* can complement the Δtr_L_ phenotype, we cloned the tracrRNA locus onto a second plasmid and introduced two T>C mutations in nucleotides 71 and 76 of tr_L_, which constitute critical residues in the −10 promoter element of P_trS_ (Fig. S2D), effectively eliminating tr_S_ expression (Fig. S2E). Expression of the double mutant, hereafter referred to as tr ^*^ restored low levels of immunity to Δtr_L_ cells (Fig. 1E, S2C), indicating that (i) Δtr_L_ is a loss-of-function allele and (ii) tr_L_ inhibits CRISPR immunity.

### tr_L_ inhibits multiple stages of CRISPR immunity

The enhanced immunity of Δtr_L_ could be due to increased rates of CRISPR adaptation, biogenesis and/or interference. We therefore sought to test the Δtr_L_ mutant in assays that measure specific stages of CRISPR immunity. To test adaptation independently of subsequent steps, we electroporated a 60 bp amplicon into cells harboring a naïve wild-type or Δtr_L_ CRISPR system. The amplicon contained a single candidate protospacer with a 5’-NGG-3’ PAM, which Cas9 recognizes during spacer selection^7^. Adaptation events were then measured by semi-quantitative PCR with a forward primer within the leader of the CRISPR array and a spacer-specific reverse primer matching the NGG-adjacent electroporated DNA sequence (Fig. 2A). Because this experiment was done in the absence of phage, the proportion of cells harboring the new spacer should solely reflect adaptation rates and not differences in phage defense resulting from biogenesis and interference. Indeed, newly acquired spacers were readily observed in Δtr_L_ cells but were below the limit of detection in cells with the wild-type CRISPR system (Fig. 2B). In parallel, we performed a second, phage-independent assay in which overexpression of the adaptation genes *cas1*, *cas2* and *csn2* enables spacer acquisition from the host genome and resident plasmids (Fig. S3A). Again, Δtr_L_ cells showed elevated adaptation rates compared to wild-type, together indicating that tr_L_ impairs spacer acquisition.

**Figure 2.**
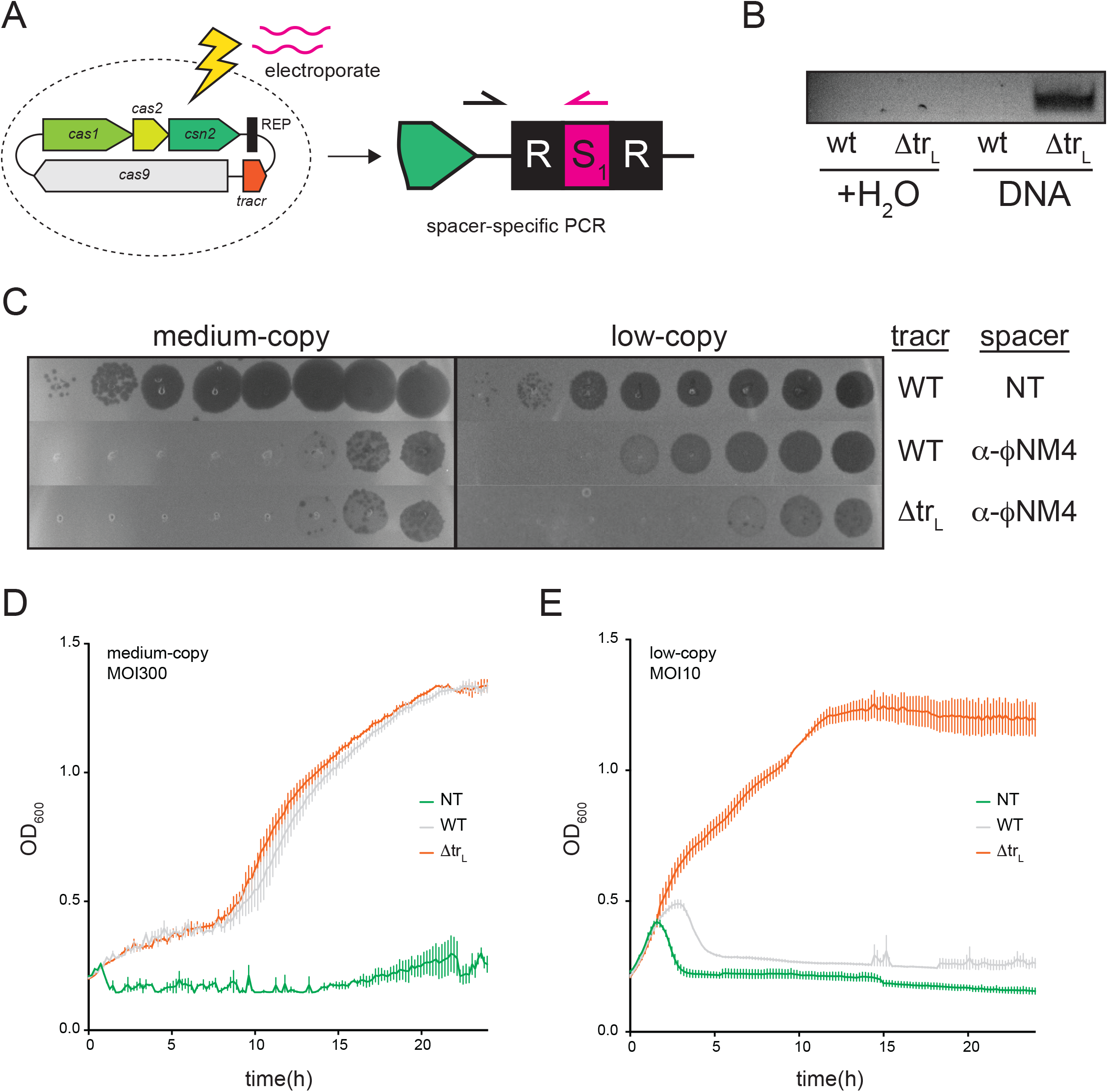
Long-form tracrRNA (tr_L_) inhibits both spacer acquisition and interference. **A)** Schematic of spacer acquisition assay. **B)** Water or a 60bp dsDNA amplicon with a single PAM was electroporated into cells with a naïve wt or ΔtrL CRISPR system, and adaptation was monitored with a spacer-specific PCR. **C)**10-fold dilutions of phage ϕNM4γ4 were plaqued on top agar lawns containing cells with medium or low-copy CRISPR plasmids harboring the indicated spacers. NT, non-targeting spacer; α-ϕNM4, spacer targeting ϕNM4γ4. **D-E)** The medium (D) and low-copy (E) strains from (C) were infected with ϕNM4γ4 in liquid culture at MOI=300 (D) or MOI=10 (E). Cell densities (OD_600_) were measured every 10 minutes in a 96-well plate reader.

To determine whether tr_L_ inhibits CRISPR biogenesis and/or interference, we programmed wild-type or Δtr_L_ CRISPR loci with a spacer targeting ϕNM4γ4 and tested phage defense using both a plaque-formation unit (PFU) assay on semi-solid agar and a liquid cell growth assay in a 96-well plate reader. On a medium-copy plasmid, the wild-type CRISPR system defends well against ϕNM4γ4, providing a 5-log reduction in PFUs and supporting bacterial growth even at an MOI of 300 (Fig. 2C-D, S3B). In this context, the Δtr_L_ mutation did not provide an added benefit. On a low-copy plasmid, the wild-type system provides only a 1-log decrease in phage PFUs and cannot support bacterial growth even at a modest MOI of 10 (Fig. 2C, 2E, S3C). Here, the Δtr_L_ mutation significantly enhances phage defense, providing an additional 2-log decrease in PFUs and supporting robust bacterial growth in liquid culture at an MOI of 10. Collectively, these results demonstrate that tr_L_ negatively regulates multiple stages of CRISPR immunity.

### tr_L_ causes widespread down-regulation of all CRISPR-Cas RNAs

tr_L_ could inhibit CRISPR immunity by affecting the levels or activity of one or more CRISPR-Cas components. To measure CRISPR-Cas expression levels, we prepared RNA from cells harboring a wild-type or Δtr_L_ CRISPR system and performed Northern blots for crRNAs and tracrRNAs, qPCR using primer pairs distributed throughout the Cas operon as well as RNAseq. In the absence of tr_L_, the levels of tr_S_, tr_P_ and processed crRNAs were all dramatically enhanced (Fig. 3A). Similarly, the protein-coding Cas genes were induced by 30 to 50-fold (Fig. 3B, S4A-C), and Cas9 protein levels showed a corresponding increase by Western blot (Fig. 3C).

**Figure 3.**
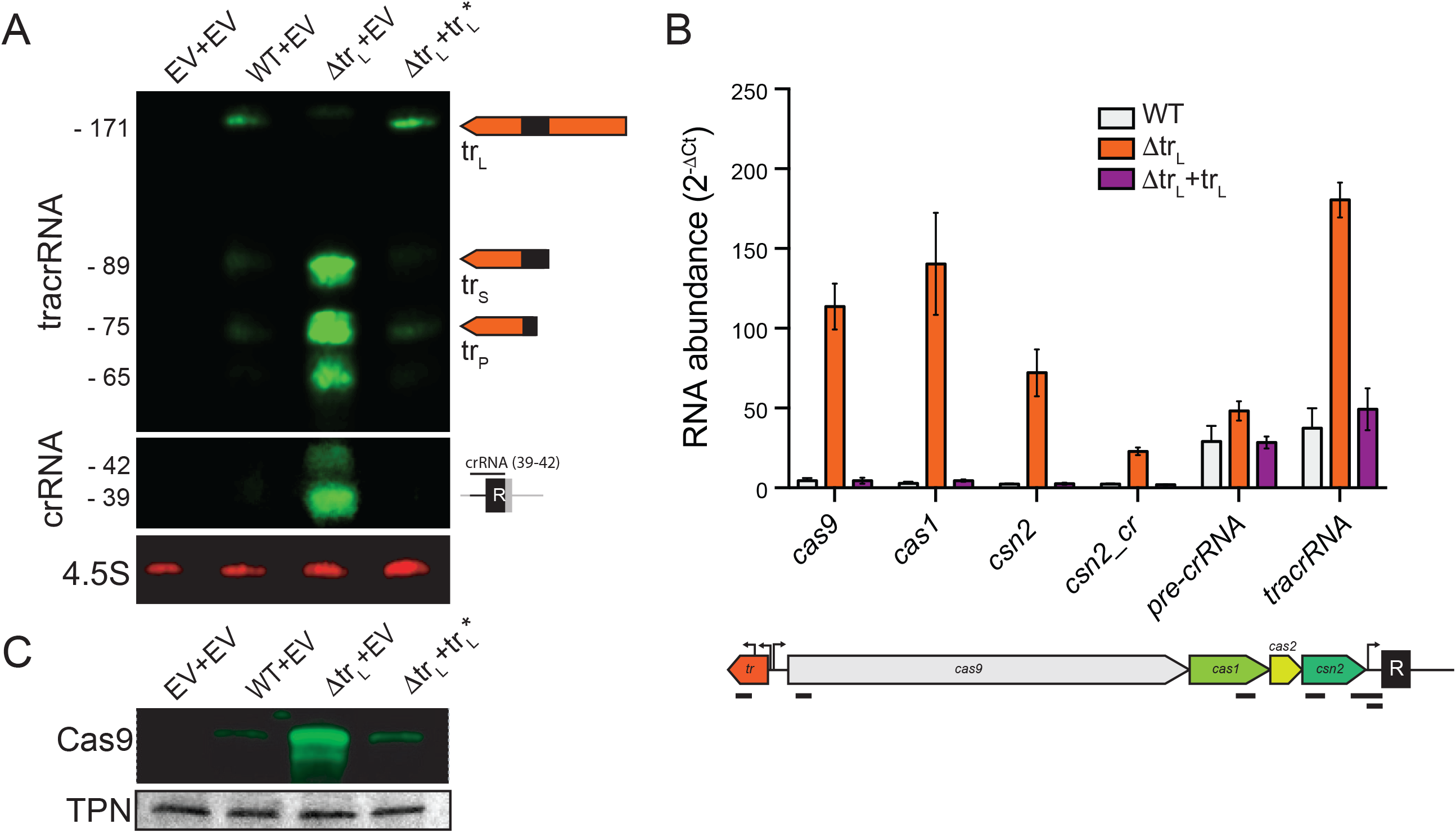
tr_L_ down-regulates all CRISPR-Cas system components. **A)** Infrared Northern blot of cells harboring the indicated plasmids grown to stationary phase. EV, empty vector; wt, naïve wild-type CRISPR system; Δtr_L_, naïve Δtr_L_ CRISPR system; tr_L_*, a plasmid expressing long-form tracrRNA. Membranes were probed with oligos matching the 3’ end of tracrRNA (top), the crRNA repeat (middle) or the 4.5S RNA (bottom, loading control). **B)** RT-qPCR of the indicated strains using primers within the coding region of *cas9*, *cas1* or *csn2*, on either side of the crRNA promoter (csn2_cr), within the pre-crRNA leader sequence or within the processed tracrRNA (tr_P_). Fold-change was expressed as 2^−ΔCt^ for each indicated primer pair relative to a *rho* control primer pair. Bottom panel shows the locations of the amplicons. **C)** Infrared Western blot of the strains in (A) using an α-Cas9 antibody. Membranes were stained with Ponceau S, and the prominent band is shown as a loading control (total protein, TPN).

Unprocessed pre-crRNA levels as measured by qPCR increased only modestly in Δtr_L_ cells, likely due to read-through from the Cas gene operon (Fig. 3B, compare ‘csn2_cr’ and ‘pre-crRNA’ primer pairs). The increase in mature crRNA levels observed by Northern blot are therefore likely due to enhanced pre-crRNA processing owing to the increased abundance of Cas9 and/or tr_S_. Collectively, these results demonstrate that tr_L_ has strong and wide-ranging inhibitory effects on the expression of all CRISPR-Cas RNAs.

### tr_L_ directs Cas9 to transcriptionally repress the Cas gene operon

The CRISPR-Cas system is transcribed from four promoters: P_trL_ and P_trS_ transcribe the tracrRNAs, P_cas9_ transcribes the Cas gene operon, and P_cr_ transcribes the CRISPR array (Fig. 1A). We next asked whether tr_L_ downregulates CRISPR-Cas RNA levels by transcriptionally repressing any of these promoters. Cells expressing Cas9 and wild-type tracrRNA or Δtr_L_ but none of the other CRISPR-Cas elements were transformed with a second plasmid harboring a transcriptional GFP fusion to each CRISPR-Cas promoter. Cas9 and tracrRNA inhibited P_cas9_ activity by roughly 50-fold but had no effect on the other promoters (Fig. 4A), consistent with the Cas gene mRNA induction observed by qRT-PCR (Fig. 3B) and RNAseq (Fig. S4B). Furthermore, these results indicated that Cas9 and tr_L_ are sufficient for repression as *cas1*, *cas2*, *csn2* and *crRNAs* were not required.

**Figure 4.**
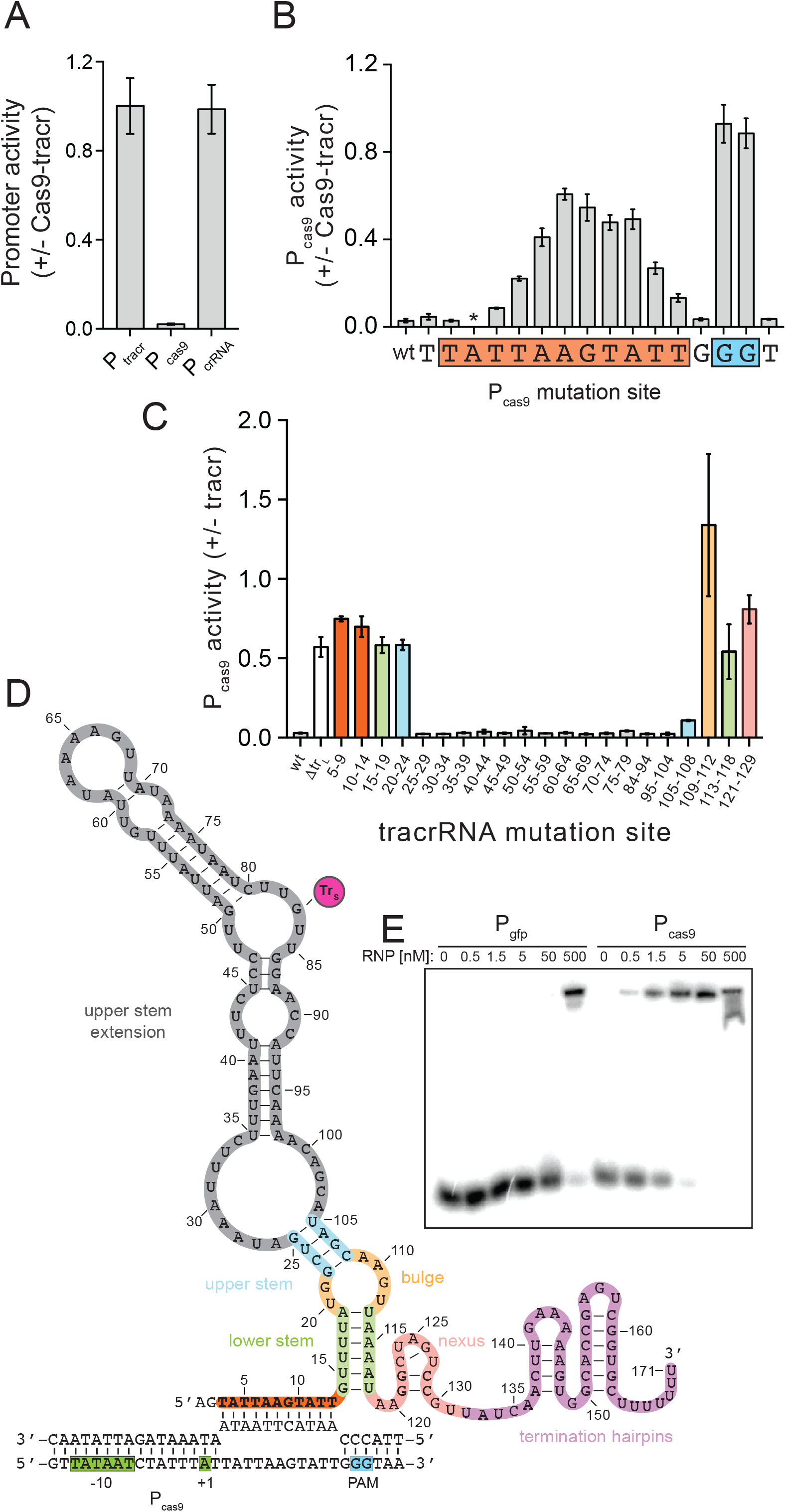
tr_L_ is a natural sgRNA (nt-sgRNA) that directs Cas9 to repress its own promoter. **A)** Promoter activity was measured (fluorescence/OD_600_) in cells harboring a plasmid expressing GFP from the indicated promoters, and a second plasmid encoding Cas9 and the full tracrRNA locus or an empty vector. **B)** Promoter activity was measured as in (A) with P_cas9_-GFP reporter constructs harboring single mutations to the complementary base at the indicated positions. The asterisk indicates that the mutation at the −10 site significantly reduced basal P_cas9_ expression (Fig. S5B), rendering the +/-Cas9-tracr comparison inconclusive. **C)** Promoter activity was measured in cells harboring a plasmid expressing Cas9 from a constitutive promoter and P_cas9_-GFP and a second plasmid expressing the indicated tracrRNA mutants or an empty vector. Sets of mutations were introduced in the tracrRNA plasmid at the indicated positions (relative to the TSS of tr_L_) to their complementary bases. **D)** Putative structure of the nt-sgRNA. Orange, P_cas9_-targeting; green, lower stem; yellow, bulge; blue, upper stem; grey, upper stem extension; pink, nexus; purple, termination hairpins; magenta circle (tr_S_), short-form tracrRNA start site. **E)** Electromobility shift assay (EMSA). A radiolabeled dsDNA derived from P_GFP_ or P_cas9_ at 50pM was mixed with Cas9:tr_L_ RNPs at the indicated concentrations at 37°C for 1 hour. Reactions were run on an 8% TBE gel.

Strikingly, inspection of P_cas9_ revealed an 11-nucleotide match between the 5’ end of tr_L_ and a region beginning 2 bp downstream from the P_cas9_ transcriptional start site (TSS) (Fig. S5A). A 5’-NGG-3’ PAM, which licenses Cas9 to bind to DNA targets specified by crRNA guides, lies immediately 3’ of the sequence match within P_cas9_. Furthermore, while Cas9 cleaves DNA targets that match the 20 spacer-derived nucleotides with a given processed crRNA, shorter match lengths of up to 16 base pairs allow stable target binding but prevent cleavage^50–52^. Our results therefore suggested that tr_L_ could direct Cas9 to bind near the P_cas9_ TSS and repress transcription while avoiding an autoimmune cleavage event.

To test this hypothesis, we individually mutated bases within the P_cas9_ 11 bp match and downstream PAM on the P_cas9_-GFP reporter plasmid (Fig. 4B, S5B). These constructs were introduced into strains harboring a second plasmid expressing Cas9 and tracrRNA or an empty vector control. tr_L_-mediated repression of P_cas9_ was abolished in both PAM mutants and greatly reduced in mutants of the 9 bases at the 3’ end of the putative 11 bp target site. These data are consistent with previous studies showing that Cas9 binding absolutely requires a PAM and strongly prefers a perfect match within the 8-12 bp “seed” region at the 3’-end of the spacer. We also tested mutations in the bases flanking the target site and PAM, as well as the ‘N’ site of the 5’-NGG-3’ PAM, which were repressed by tr_L_ at wild-type levels. To confirm that tr_L_-mediated repression of P_cas9_ requires Cas9 protein, we combined a plasmid expressing tracrRNA and a *cas9* nonsense mutant with the P_cas9_-GFP reporter. Indeed, tr_L_-mediated P_cas9_ repression required Cas9 (Fig. S5C) although tracrRNAs were undetectable in *cas9* null mutants (Fig. S5D), likely because they are protected from degradation by Cas9 protein. Together, our results are consistent with a model in which the 5’ end of tr_L_ directs Cas9 to bind its own promoter and repress transcription of the Cas gene operon.

### tr_L_ is a natural single guide RNA

To comprehensively probe the determinants of tr_L_ repression, we generated 5 nt mutations tiled throughout the tracrRNA region specific to tr_L_ as well as variably sized mutations in tracrRNA domains within tr_S_ with well characterized functions, including the upper stem, bulge, lower stem and nexus (Fig. 4C, S6D). Each tracrRNA mutant was introduced on a plasmid into cells harboring a second plasmid expressing both *cas9* from a constitutive promoter and a P_cas9_-GFP reporter. As expected, mutations in the nexus and bulge, tracrRNA regions critical for Cas9 binding and targeting, alleviated tr_L_ repression of P_cas9_. Mutations in the 11 nt P_cas9_-matching sequence similarly blocked repression, confirming the ability of this region to guide tr_L_ to its target. Curiously, mutations in the lower stem also blocked tr_L_ repression, while mutations in the upper stem had a milder but significant effect. The tracrRNA upper and lower stem regions normally base pair with complementary sites in the pre-crRNA repeats, enabling processing of the RNA duplex (Fig. S6B-C). However, our previous findings demonstrated that tr_L_-mediated repression of P_cas9_ occurred in the absence of crRNA (Fig. 4B), indicating that the upper and lower stems of tr_L_ are not occupied by their cognate crRNA residues in the Cas9:tr_L_ targeting complex. Instead, we noticed that the 12 nucleotides just downstream from the P_cas9_-targeting region of tr_L_ almost perfectly mimic those at the 5’ end of the crRNA repeat, suggesting that tr_L_ could fold in on itself to reconstitute the upper and lower stems (Fig. 4D). Indeed, the crRNA mimicking residues were also critical for repression (Fig. 4C), suggesting that in the absence of crRNAs, tr_L_ could form a natural single-guide RNA in which the P_cas9_-targeting region takes the place of the crRNA spacer and the upper and lower stems are formed intramolecularly (Fig 4D, S6A).

In the putative natural single guide structure, tr_L_ residues 14-19 and 113-118 base pair to form the lower stem, and 5 nt mutations in either strand that would prevent base pairing also disrupt tr_L_ repression (Fig. 4C, green bars, 4D). To interrogate this structure, we generated a construct in which both lower stem strands are simultaneously mutated to their complement, which should restore base pairing potential. Prior studies on sgRNA activity demonstrated that sequence changes within the lower stem are tolerated as long as base-pairing is maintained^53^. Indeed, high levels of repression were observed in the double mutant (Fig. S5E-F), supporting our model that tr_L_ residues 14-19 mimic the crRNA repeat and form the lower stem. Another feature of the natural single guide is the large 79 nt “upper stem extension” which is replaced by an 8 nt duplex or 5’-GAAA-3’ tetraloop in the crRNA and synthetic sgRNA respectively (Fig. S6A-D). To understand whether this region is dispensable for tr_L_ repression, we replaced the entire 79 nt segment with a 5’-GAAA-3’ tetraloop and found that repression of P_cas9_ was maintained (Fig. S5E). Finally, to test whether Cas9 and tr_L_ are sufficient to form a repressive complex on P_cas9_, we performed an EMSA assay with purified Cas9, *in vitro* transcribed tr_L_ and a 55 bp dsDNA derived from the tr_L_-targeted within P_cas9_. Indeed, Cas9:tr_L_ bound P_cas9_ with an affinity of roughly 1 nM, which was considerably tighter than the nonspecific affinity of Cas9:tr_L_ for a dsDNA derived from a non-targeted control promoter with a single PAM (Fig. 4E).

To confirm that tr_L_ targeting is sequence-specific, owing to base-pairing within the 11 nt P_cas9_-matching region, we reprogrammed tr_L_ to target two new 11 bp sites within promoters unrelated to P_cas9_ on GFP reporter plasmids (Fig. 5A). In each case, promoter activity was significantly reduced in the presence of the reprogrammed constructs but not the native tr_L_. Next, we tested the effects of varying the tr_L_ match length and found that increasing complementarity from 11 to 13 bp resulted in an additional 2-fold decrease in promoter activity (Fig. S7A). While a 15 bp match did not further reduce promoter activity, we could not readily transform cells harboring tr_L_ constructs with GFP reporter plasmids containing a >16 bp tr_L_ match (Fig. S7B). Previous studies showed that sgRNAs with match lengths greater than 16 bp enable Cas9 to cleave its target^50–52^, suggesting that the low transformation efficiencies we observed could be due to cleavage mediated by reprogrammed tr_L_. To explore this possibility, we performed an *in vitro* Cas9 cleavage assay with tr_L_ constructs harboring increasing match lengths to a 1 kb amplicon derived from the GFP reporter plasmid pCN57. Cleavage was first apparent with a 17 nt match, and with a 19 nt match, cleavage activity resembled that of *bona fide* sgRNAs (Fig. 5B). Collectively, our results suggest that tr_L_ evolved as a natural single guide to strongly repress, but not cleave, P_cas9_ and maintain low-levels of Cas operon expression.

**Figure 5.**
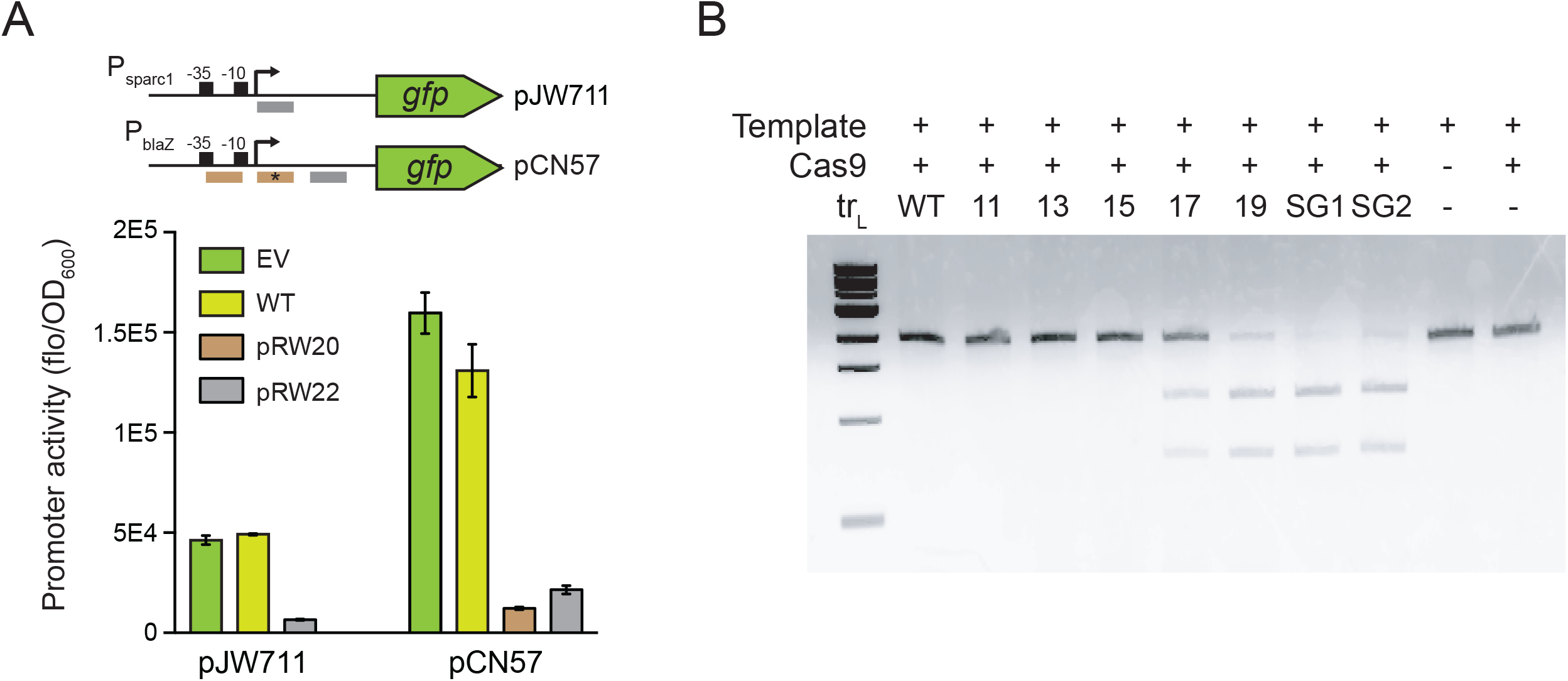
tr_L_ can be reprogrammed to direct Cas9 repression or cleavage of novel targets. **A)** Promoter activity was measured in cells harboring the GFP reporter plasmid pJW711 or pCN57 and a second plasmid expressing Cas9 and the indicated tr_L_ variants. EV, empty vector; WT, the wild-type P_cas9_-targeting tr_L_; pRW20 and pRW22, tr_L_ reprogrammed to target the 11bp regions indicated in brown or gray bars respectively. The asterisk indicates a second pRW20 binding site within pCN57 containing a single mismatch at the −9 seed position relative to the PAM. **B)***In vitro* cleavage assay. 30 nM sgRNA or tr_L_, 33.3 nM Cas9 and a 3 nM 1kb amplicon from pCN57 were incubated at RT for 10 min and cleavage products were run on a 1.5% agarose gel. WT, tr_L_; 11-19, tr_L_ variants transcribed from pRW22 with an expanded pCN57 targeting sequence of the indicated length; SG1-2, replicates of an sgRNA with a 20nt pCN57 targeting sequence.

### tr_L_ controls a system-wide switch that is responsive to crRNA expression

We next investigated whether the immunosuppressive effects of tr_L_ were solely due to repression of P_cas9_, or whether tr_L_ could also inhibit other CRISPR functions, for instance by preventing Cas9 and/or crRNAs from performing their canonical roles in adaptation, biogenesis, or interference. We constructed a plasmid encoding a naïve CRISPR-Cas system with a PAM mutation (P_cas9_^NGC^) that renders P_cas9_ insensitive to tr_L_ repression (Fig. 4A). We then tested the P_cas9_^NGC^ strain in a top agar immunity assay and found high levels of phage resistance comparable to the Δtr_L_ mutant (Fig 6A), indicating that transcriptional repression is the primary if not sole inhibitory function of tr_L_. As expected, despite the presence of tr_L_, P_cas9_^NGC^ cells overexpress Cas9 at Δtr_L_ levels (Fig. S8A). Interestingly, this construct also phenocopies Δtr_L_ with high levels of tr_S_, tr_P_, and crRNA, none of which are transcribed from P_cas9_ (Fig. S8A). Given that tracrRNAs and crRNAs were undetectable in the absence of Cas9 (Fig. S5D), we wondered whether their abundance in Δtr_L_ and P_cas9_^NGC^ could stem from Cas9 binding and stabilization. To test this hypothesis, we overexpressed Cas9 in cells harboring a wild-type CRISPR system and found that tr_S_, tr_P_, and crRNAs were all significantly upregulated (Fig. S8B). These results suggest that tr_L_ serves as a master regulator for the entire CRISPR-Cas system by (i) directly controlling the protein-coding Cas operon through P_cas9_ and (ii) indirectly controlling tracrRNA and crRNA levels by regulating Cas9 abundance.

**Figure 6.**
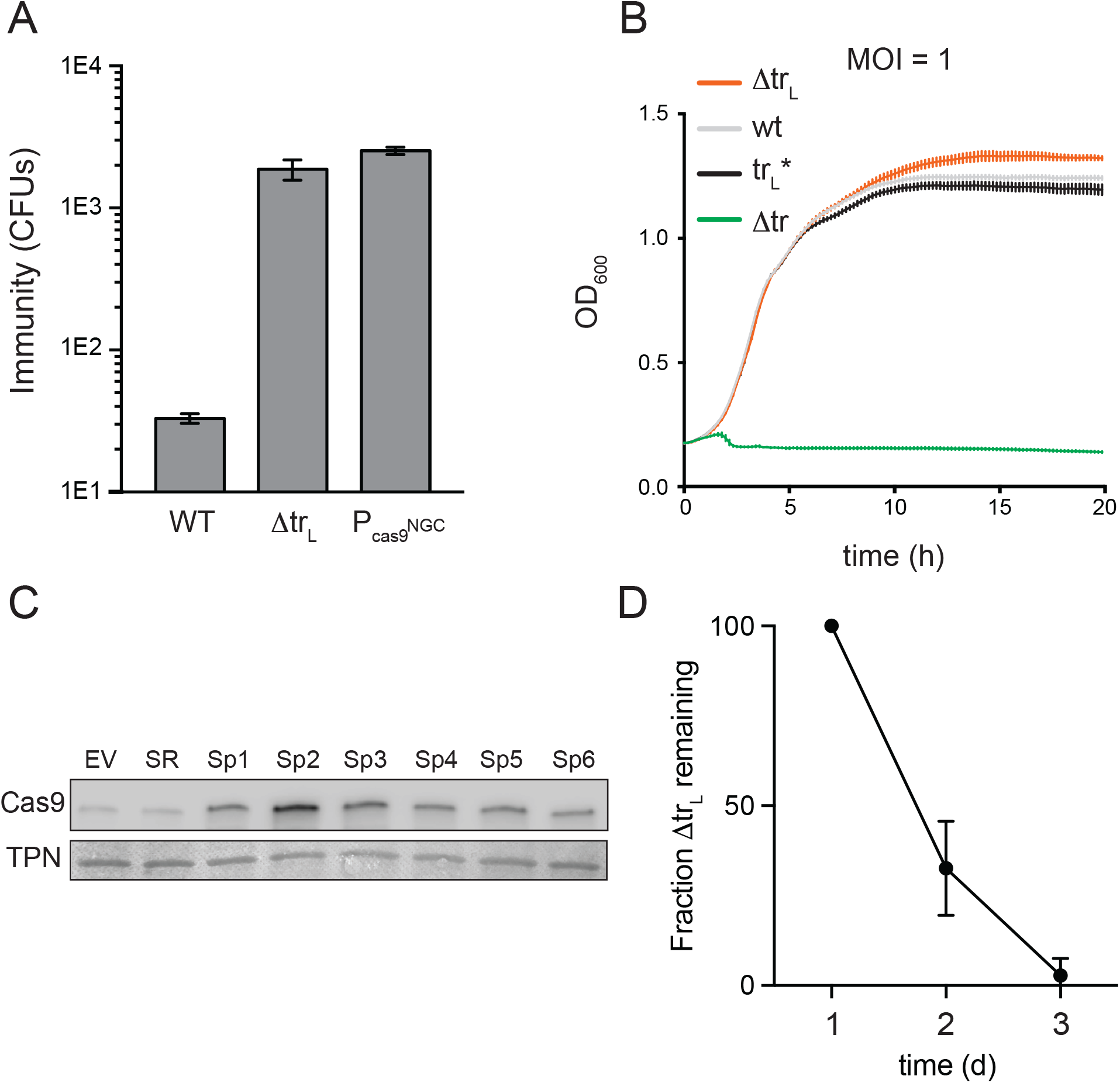
tr_L_ is master switch that controls auto-immunity and responds to crRNA spacer identity. **A)** Naïve CRISPR-Cas mutants were infected with ϕNM4γ4 at MOI = 25 in top agar, and surviving colonies were quantified. Δtr_L_, deletion of the long-form tracrRNA; P_cas9_^NGC^, PAM mutation in the tr_L_ target site within P_cas9_. **B)** Interference assay with ϕNM4γ4 at MOI = 1 with cells harboring a plasmid expressing Cas9, the ϕNM4γ4-targeting spacer NM2, and the indicated tracrRNA mutants. tr_L_*, long-form tracrRNA only; Δtr, full deletion of the tracrRNA locus. The tr_L_* strain harbored a second plasmid expressing additional tr_L_* from P_sparc2_ in order to ensure levels of tr_L_ expression comparable to tr_S_ in the Δtr_L_ strain. All other strains harbored a second empty vector. IPTG was added at 1 mM to induce expression from P_sparc2_. **C)** Chemiluminescent Western blot using an α-Cas9 antibody on logarithmic phase cells harboring a plasmid expressing the wild-type CRISPR system with the CRISPR array deleted and a second plasmid expressing no CRISPR array (EV) a single repeat (SR) or a single spacer from the endogenous *S. pyogenes* CRISPR array (Sp1-6). Quantification of biological triplicates shown in Fig. S8F. **D)** Cells harboring a plasmid expressing a naïve wild-type or Δtr_L_ CRISPR system were co-cultured in a competition assay. Cells were grown to stationary phase and diluted into logarithmic phase twice per day, and each morning, cultures were plated and wild-type and Δtr_L_ cells were counted by PCR of the tracrRNA locus.

We next asked whether tr_L_ is a specialized transcriptional repressor or whether it can also bind crRNAs like tr_S_. To explore this possibility, we performed interference assays on cells harboring CRISPR systems expressing either tr_L_ (tr_L_^*^) or trS (Δtr_L_), each programmed with the ϕNM4γ4-targeting spacer NM2. Both systems were engineered to include the P_cas9_^NGC^ mutation in order to normalize Cas gene expression levels across experiments. In a top agar interference assay (Fig. S8C), or a liquid interference assay at low MOI (Fig. 6B), tr_L_ and tr_S_ provided similar levels of protection against ϕNM4γ4. In liquid interference assays at high MOI (Fig. S8D), tr_L_ provided less protection than tr_S_, possibly because tr_L_ is processed poorly (Fig. S8E), forming less of the mature targeting complex Cas9:tr_P_:crRNA. Nonetheless, these results indicate that tr_L_ can hybridize with crRNAs in order to interfere with viral targets specified by the crRNA spacer.

Once bound, crRNAs occupy the upper and lower stems of tr_L_ preventing formation of the natural single guide (Fig. 4D, S6E). We therefore wondered whether crRNA expression interferes with tr_L_-mediated repression of P_cas9_. To explore this possibility, we measured Cas9 levels by Western blot in cells harboring a CRISPR-Cas system with no CRISPR array, a single repeat or an individual spacer from the natural *S. pyogenes* SF370 CRISPR array. We observed low Cas9 levels in cells without a CRISPR array, consistent with unimpeded P_cas9_ repression by Cas9:tr_L_ (Fig. 6C, S8F). The presence of a single repeat did not enhance Cas9 expression, indicating that a naïve pre-crRNA did not appreciably interfere with tr_L_ repression; however, the presence of any single spacer caused up-regulation of Cas9 (Fig. 6C, S8F). These results suggest the presence of a regulatory circuit in which crRNAs provide feedback through tr_L_ to affect P_cas9_ expression. Notably, the CRISPR array is transcribed from its own promoter providing another entry point for system-wide regulation.

### CRISPR-Cas repression by trL inhibits auto-immunity

Our results indicate that in wild-type cells, tr_L_ maintains the CRISPR-Cas system in a lowly active state. Given that inactivation of tr_L_ leads to enhanced immunity, why does the system encode a P_cas9_ repressor? One possibility is that constitutive expression of CRISPR-Cas components could cause autoimmunity, stemming from off-target adaptation or interference against the bacterial chromosome or resident plasmids^44–47,54,55^. To gauge whether cells harboring a de-repressed CRISPR system suffer a viability defect, we performed a liquid growth competition experiment. Cells harboring a wild-type or Δtr_L_ CRISPR system were mixed in equal parts and serially passaged twice a day. Each day, aliquots were plated to single colonies and the relative proportions of wild-type and Δtr_L_ cells were determined by PCR. After one and two days, the number of Δtr_L_ cells dropped to 33% and 3% of their original number respectively (Fig. 6D), indicating that they suffer a growth defect compared to cells with a wild-type CRISPR system. Together, our results suggest that tr_L_ could repress the CRISPR system to avoid autoimmunity while allowing enough expression for some level of viral surveillance (Fig. 2B-E).

### TracrRNA regulation is dynamic on evolutionary timescales

We next investigated whether tr_L_-mediated repression of P_cas9_ is specific to *S. pyogenes* or conserved in other type II-A CRISPR-Cas systems. In a list of previously annotated short-form tracrRNA loci^56^, we examined ‘branch 1’ which contains 80 representatives from *Streptococci*, *Listeria* and *Lactobacilli*. We queried the presence of tr_L_ regulation by looking for 3 criteria: (i) an 11 bp or greater match between a putative tr_L_ and a region predicted to contain the Cas9 promoter, (ii) a 6 bp lower stem sequence (5’-GTTTTA-3’) just downstream from the tr_L_ targeting site and (iii) a PAM sequence (5’-NGG-3’) just downstream from the P_cas9_ target site. CRISPR systems that fit all three criteria, allowing for a single mismatch, were designated as “tr_L_^+^” and the others as “tr_L_^−^”. Of the 80 CRISPR loci surveyed, 34 (43%) are tr_L_^+^ including one strain with a 10 bp match, two with a 5’-GTCTTA-3’ lower stem and one with a 5’-NAG-3’ PAM (Fig. 7A, Table S2). tr_L_^+^ genomes were not constrained to specific phylogenetic clusters but were distributed intermittently throughout the tree, suggesting frequent loss or gain events. Curiously, while the lower stems and PAMs showed near perfect conservation among tr_L_^+^ loci, the identity of the matching sequence varied and its length ranged from 11-15 bp, indicating covariation over evolutionary timescales between the tr_L_ and P_cas9_ targeting determinants (Fig. 7A, S9). Variability was also observed in the length and sequence of the tr_L_ upper stem extension, the precise location of the targeted sequence relative to *cas9* promoter elements, and the location of P_cas9_ relative to the *tracrRNA* or *cas9* coding regions (Fig. S10-11). Within tr_L_^−^loci, the intergenic space between tr_S_ and *cas9* was on average roughly 200 bp shorter than in tr_L_^+^ loci (Fig. 7B) owing to deletions that removed the targeting and/or targeted sites (Fig. S11B). In other cases, we observed tr_L_^−^loci with intergenic lengths comparable to tr_L_^+^ that contained one or more mismatches within the seed but retained perfect PAMs and lower stems (Fig. S11C). Our data suggests that similar seed mutants can retain intermediate levels of repression (Fig. 4B), while deletions of targeting determinants likely result in complete de-repression (Fig. 4B). Collectively, our results demonstrate that tr_L_-mediated regulation of P_cas9_ is common in type II CRISPR-Cas systems. We speculate that individual strains have utilized, jettisoned or mutated this regulatory module in response to changes in phage predation associated with exploration of new hosts and environments.

**Figure 7.**
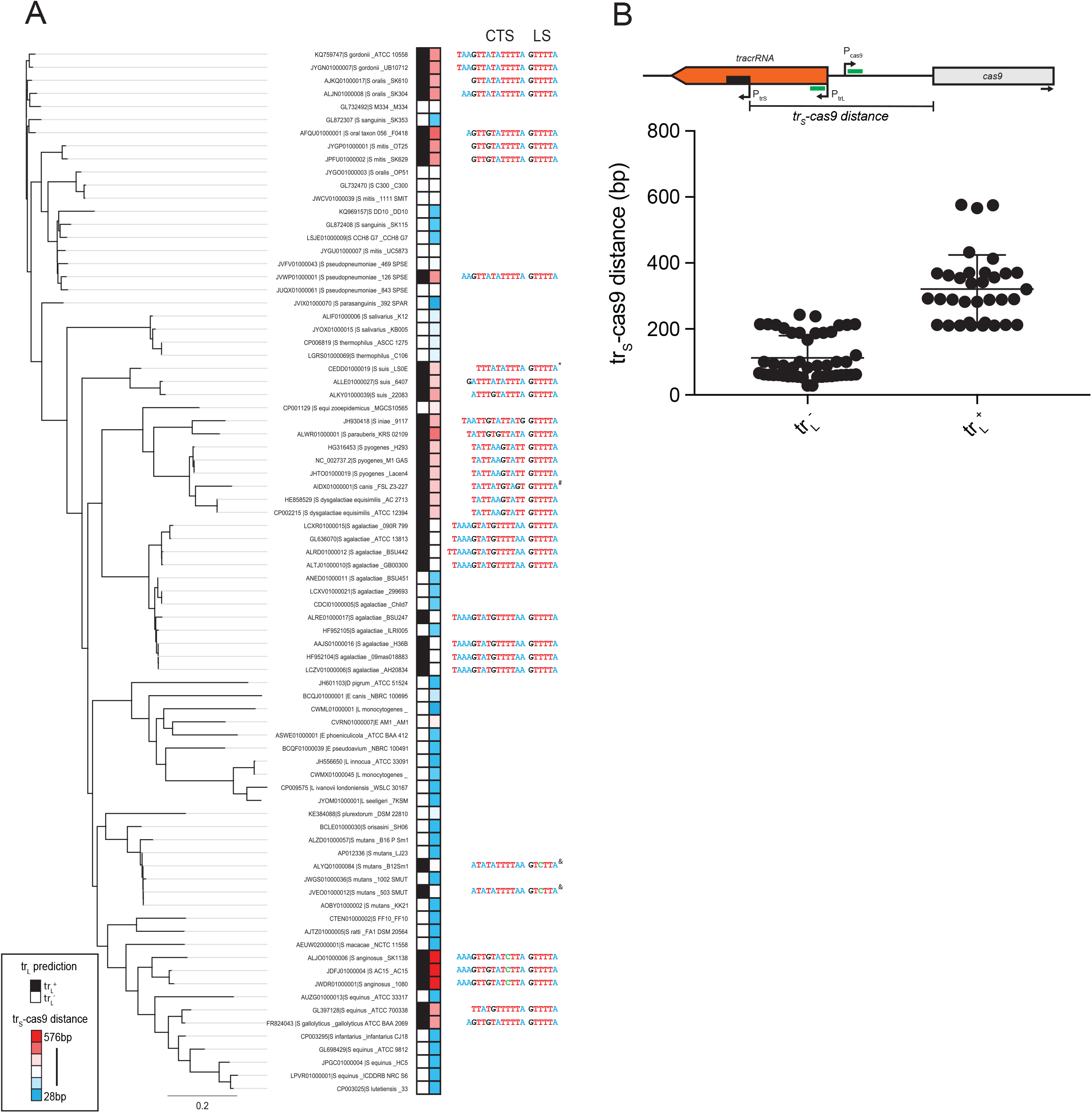
tr_L_ is a conserved and dynamic feature of type II-A CRISPR-Cas systems. **A)** A phylogenetic tree was constructed using Cas9 protein sequences from representative type II-A ‘branch-1’ genomes. Each CRISPR-Cas locus was queried for the presence of tr_L_-mediated regulation of P_cas9_ (black boxes, tr_L_^+^; white boxes, tr_L_^−^) by looking for an 11 bp candidate targeting sequence (CTS), a 5’-GTTTTA-3’ lower stem (LS) and target site 5’-NGG-3’ PAM. The intergenic distance between the translation start site of Cas9 and the transcriptional start site of the short-form tracrRNA (tr_S_-cas9) was measured and plotted as a heatmap (red boxes = longer distances, blue boxes = shorter distances). The CTS and lower stem LS are listed for each tr_L_^+^ locus, with nucleotides colored by identity. *, 10 bp CTS; #, 5’-NAG-3’ PAM; &, 5’-GTCTTA-3’ LS; scale bar, average amino acid substitutions per site. Full metadata available in Table S2. **B)** Schematic showing the intergenic distance between tr_S_ and Cas9 (tr_S_-cas9, black bar) with putative targeting determinants (green bars). Promoters are shown with arrows indicating the transcriptional start sites. Below, intergenic distances were plotted for tr_L_^+^ and tr_L_^−^loci.

## DISCUSSION

How CRISPR-Cas systems are regulated to enhance the targeting of foreign nucleic acids while avoiding autoimmunity is a fundamental yet poorly understood aspect of CRISPR biology. In the absence of dedicated Cas-encoded transcription factors, a mechanistic understanding of whether and how type II CRISPR-Cas9 systems are regulated was unknown. Here, we report that long-form tracrRNAs are capable of forming natural single guides that direct Cas9 to transcriptionally repress the Cas operon (Fig. 4A, S6, summarized in Fig. 8). As a result, the CRISPR-Cas system is kept in a lowly active state that is sufficient to interfere against remembered threats (Fig. 2C-E) but adapts poorly to unrecognized phages. De-repression of the system, by genetic ablation of tr_L_, results in a roughly 50-fold increase in CRISPR-Cas components (Fig. 3, S4) and a 3,000-fold increase in immunization rates against an unrecognized phage (Fig. 1B-E). As with many immune systems, this hyperactivity comes at a price; cells with de-repressed systems are outcompeted by wild-type cells in the absence of a phage threat (Fig. 6D). Below, we propose that bacteria navigate these costs and benefits through (i) mutation of the regulatory controls and (ii) transient de-repression of P_cas9_:tr_L_.

**Figure 8.**
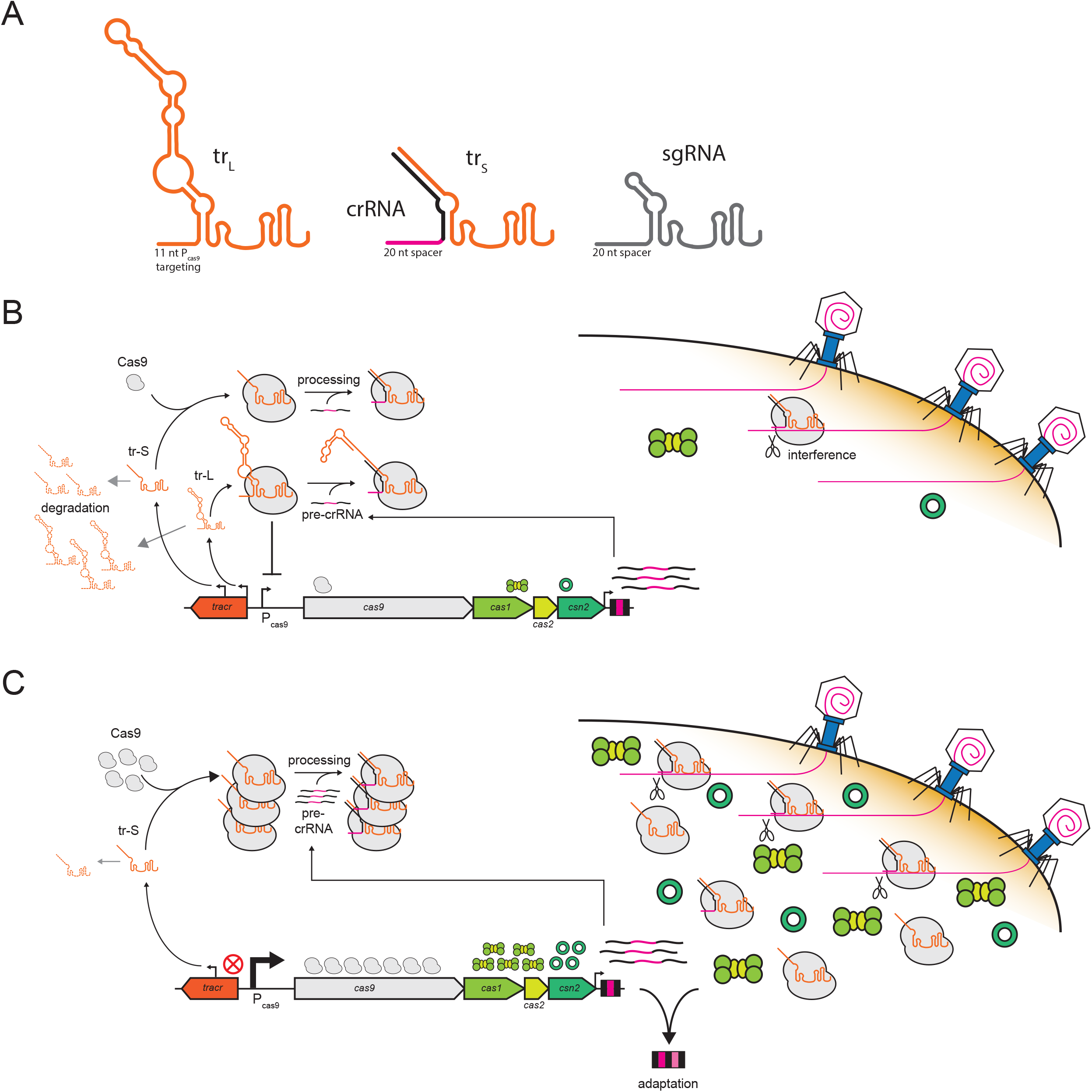
Model of tr_L_ regulation. **A)** tr_L_ forms a natural single guide RNA that preserves the structural elements of the tracr:crRNA duplex and artificial sgRNAs. **B)** In wild-type cells, trL represses P_cas9_ resulting in low levels of Cas gene expression. Low Cas9 levels destabilize tracrRNAs and lead to low levels of tracrRNA:crRNA processing. In this state, the CRISPR-Cas system can interfere against viruses at low MOIs but cannot effectively acquire new spacers. crRNAs partially relieve repression by binding to tr_L_ and preventing formation of the natural single guide. **C)** In the absence of tr_L_, Cas gene expression is induced, tr_S_ is stabilized by Cas9 and tracrRNA:crRNA processing is enhanced. As a result, interference can occur at higher MOIs and spacer acquisition is stimulated.

Using a bioinformatic approach, we identified tr_L_ transcriptional regulators throughout many branches of the type II-A CRISPR-Cas tree (Fig. 7A). Moreover, we found evidence that the regulatory elements are frequently lost, suggesting that individual strains can use or lose this system according to their unique needs. In cases where the regulator is wholly absent, high basal rates of CRISPR-Cas expression could enable survival in environments with abundant phage. It is unclear whether and how these strains manage the autoimmunity associated with tr_L_ loss. In other cases, mutations in the seed, lower stem or PAM could only partially relieve repression resulting in slightly elevated rates of basal CRISPR-Cas expression (Fig. 4B). These strains may better survive phage-replete environments while mitigating the autoimmunity associated with full tr_L_ loss. Furthermore, we speculate that seed mutants in particular could represent evolutionary stepping stones, where a second compensatory mutation could restore repression, resulting in the covariation we observed between the targeting and targeted sites throughout the type II-A tr_L_ tree. By introducing a single mutation at either site, individuals within a bacterial community can sample modifications to P_cas9_, affecting basal expression levels and/or regulatory strength, to prepare the community for changing environments. Work in our lab seeks to understand whether tr_L_-regulation extends to more distant branches of the type II tree, and whether tr_L_ can target other promoters, either within or beyond the CRISPR locus. In the *F. novicida* type II-B CRISPR system, tracrRNA hybridizes with a non-canonical ‘scaRNA’ that utilizes an 11 bp targeting sequence to transcriptionally repress an immunostimulatory lipoprotein^52^ in the bacterial genome. Our work shows that scaRNAs are not required for Cas9-mediated transcriptional control, greatly expanding the list of CRISPR systems with candidate regulators. Further, we show that transcriptional regulation by Cas9 is not restricted to *F. novicida* but may be a conserved property across type II CRISPR-Cas systems.

We expect that Cas9:tr_L_ repression can also be transiently relieved in response to an unknown stimulus. We note that the configuration of the *S. pyogenes* CRISPR-Cas regulatory circuit offers several entry points for transient induction. Transcriptional changes in the activity of P_trL_, P_trS_ or P_cr_ could each induce P_cas9_ by decreasing the levels or repressive potential of tr_L_. In the simplest case, inhibition of P_trL_ would alleviate P_cas9_ repression by downregulating tr_L_ itself. P_trS_ activation could destabilize tr_L_ as it is outcompeted by tr_S_ for Cas9 binding and protection. P_cr_ induction could inactivate tr_L_ as the upper and lower stems are bound by accumulating pre-crRNAs, preventing folding of the natural single guide. Processing of tr_L_ by RNAseIII separates the 5’ targeting region from the 3’ nexus and termination hairpins, which are required for Cas9 binding. Therefore, cellular conditions that upregulate or activate RNAseIII could transiently induce CRISPR-Cas expression. Another intriguing candidate for post-transcriptional regulation is the 79 nt upper stem extension of the natural single guide, which is wholly dispensable for repression (Fig. 4C) and likely to be extruded from the main body of Cas9^57^. Secondary structural predictions of this region include a series of hairpins and loops (Fig. 4D)^58^ which could serve as substrates for cleavage, sequestration or re-folding by host proteins and/or small molecules.

Another candidate mechanism for transient CRISPR-Cas induction is the sequestration of Cas9:tr_L_ to a crRNA-targeted phage during an infection. In archaeal type I-A systems, the Cas-encoded transcription factor Csa3b associates with the Cascade interference complex, and the introduction of a viral target de-represses CRISPR-Cas expression by sequestering Cascade:Csa3b. It remains unclear whether this induction is necessary for viral defense or if a similar strategy could provide protection in bacteria given the short lifecycle of a bacteriophage compared to an archaeal virus. The signals and mechanisms of transient induction, especially those occurring in the native *S. pyogenes* host, are active areas of exploration in our lab.

Our results provide the first example of an intrinsic CRISPR-Cas regulator within a bacterial host. The emerging literature on CRISPR-Cas regulation in bacteria focuses on transcription factors or chaperones encoded in the genomes of hosts with type I CRISPR-systems^20^. In these cases, CRISPR-Cas promoters likely evolved to join existing regulons, enabling CRISPR activities to be tied to a variety of internal and environmental stimuli. Less clear is how and whether CRISPR-Cas systems are regulated following horizontal transfer into a new host, a major route of CRISPR-Cas evolution, particularly for type II systems which are present in diverse animal-associated microbiomes^59^. We believe that intrinsic regulators like tr_L_ could facilitate horizontal transfer by dampening autoimmunity upon delivery and allowing the new bacterial host to tune CRISPR-Cas9 expression to meet its needs, through mutation or deletion of the control elements. Whether type II systems join host regulons over time, use only intrinsic control, or evolve overlapping extrinsic and intrinsic pathways are open questions. Archaeal type I-A systems are the lone other example of intrinsic regulation, mediated by a Cas-encoded transcription factor^25^. Our results show that intrinsic control does not require dedicated transcription factors and may be a more general property of CRISPR-Cas biology. It remains to be seen whether other CRISPR-Cas types have evolved their own RNA-guided intrinsic regulators, for instance using degenerate self-targeting spacers^60^.

Discoveries in basic CRISPR-Cas biology have continually led to new rounds of CRISPR tool development. We believe a better understanding of how tr_L_ is naturally regulated will facilitate the development of controllable single guides. The recent creation of a theophylline-responsive Cas9 RNP by the fusion of an aptamer to the sgRNA upper stem^61^ underscores the potential for regulation to occur through the solvent-exposed 79 nt upper stem extension of tr_L_. Furthermore, an elucidation of the structure and function of the Cas9:tr_L_:dsDNA repressive complex could inform a new generation of Cas9-based transcriptional tools. As the list of sequenced microbial genomes grows, so does the catalog of novel Cas9 orthologs, each with the possibility of new PAMs and unique cleavage activities. Our results suggest that many of these genomes might be worth a closer look for the blueprints of natural single guides.

## ACKNOWLEDGEMENTS

We would like to thank Michael Laub, Luciano Marraffini, Geraldine Seydoux and Samuel Sternberg for their comments on the manuscript. We thank Karole D’Orazio and members of Rachel Green’s lab for sharing technical assistance and reagents for *in vitro* experiments. We thank Jeremy Nathans’ and Carol Greider’s lab for sharing reagents and equipment. We thank David Mohr and the GRCF High Throughput Sequencing Center for assistance with NGS experiments.

## AUTHOR CONTRIBUTIONS

R.E.W., T.P., B.T.K.N., L.W.G. and J.W.M. designed and executed the research studies. E.S., S.M.S., and M.J.S. assisted with plasmid construction. R.E.W. and J.W.M. wrote the manuscript.

---

**Figure S1.**
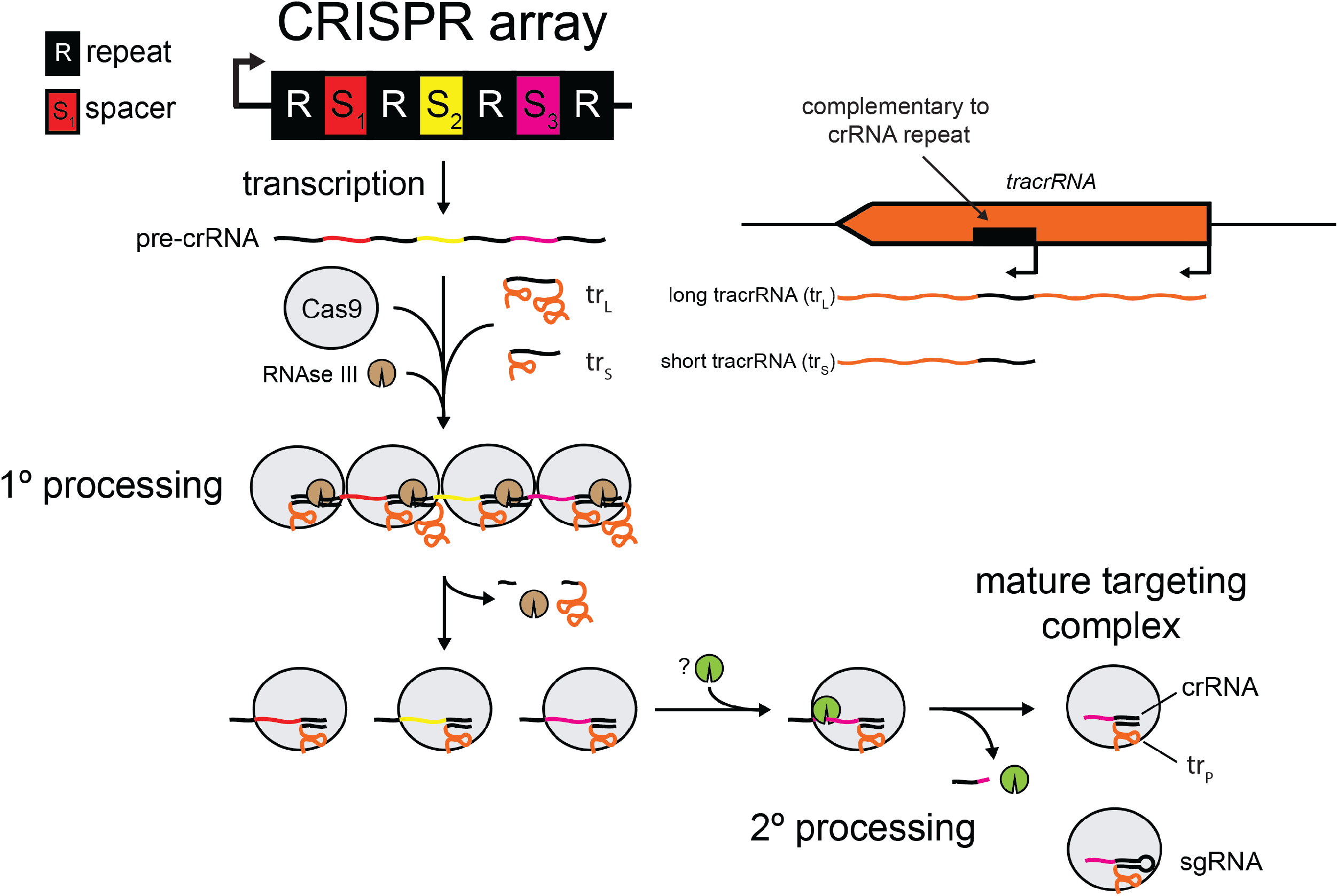
Overview of CRISPR biogenesis. The CRISPR array is transcribed as one long precursor CRISPR RNA (pre-crRNA). tracrRNAs are transcribed from one of two promoters, producing a long-form (tr_L_) and short-form (tr_S_). Once bound to Cas9, tr_S_ or tr_L_ can recruit the pre-crRNA through base-pairing between the pre-crRNA repeats (black lines) and a complementary region within tr_S_/tr_L_ (black line). RNAseIII, a bacterial enzyme that cleaves dsRNAs, cuts the tracrRNA:pre-crRNA duplex in the middle of the repeat (1° processing, Fig. S6), releasing RNAseIII and the cleaved 5’ end of tr_S_/tr_L_. Following cleavage between all crRNA repeats, Cas9 and the processed trRNA (tr_P_) remain bound to a pre-crRNA fragment containing a single 30nt spacer with half repeats on either side. Next, a second processing event digests the 5’ end of the pre-crRNA fragment leaving a 20nt spacer (2° processing) and producing the mature targeting complex (Cas9:tr_P_:crRNA).

**Figure S2.**
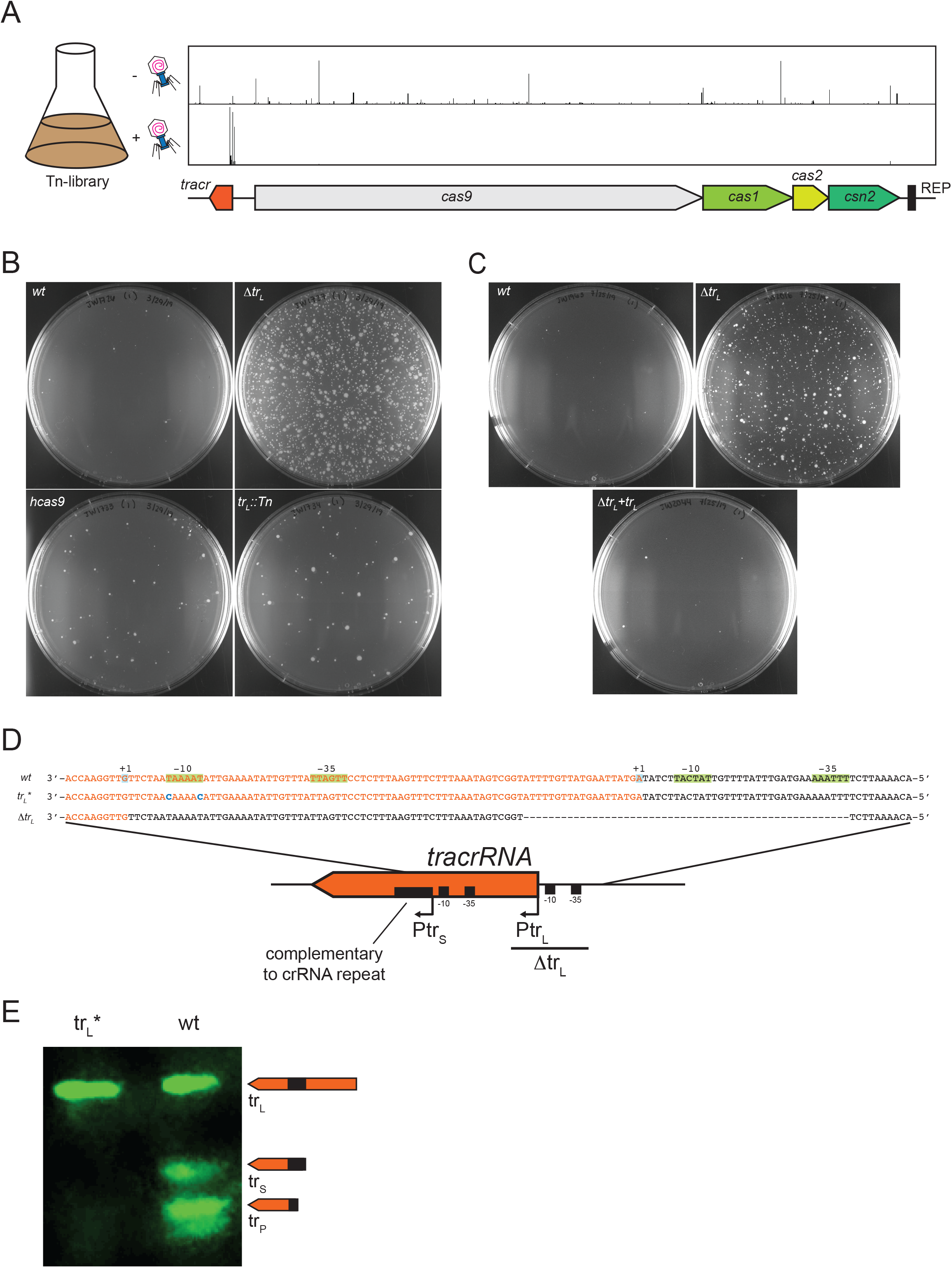
Characterization of tracrRNA mutants. **A)** An extended view of Tn-seq reads in the CRISPR-Cas region for representative – phage (top panel) and + phage (bottom panel) experiments. The location of each hit corresponds to the gene annotations shown in cartoon format below the Tn-seq reads. **B-C)** Representative plates are shown from the top agar immunity assays performed in Fig. 1D-E. **D)** Schematic of the tracrRNA promoter region. The wild-type sequence is shown in the top row with transcribed sequences shown in orange. P_trL_ and P_trS_ are each indicated with a transcriptional start site (+1) and the −10 and −35 promoter elements that contact RNA polymerase. In tr_L_^*^ (middle row), two T>C mutations in the tr_S_ −10 element were introduced to prevent tr_S_ transcription. In Δtr_L_ (bottom row), the tr_L_ promoter and 19 5’ nucleotides have been deleted. **E)** Infrared northern blot on cells harboring a plasmid expressing a naïve wild-type or tr_L_* CRISPR-Cas system probed with oligos matching the 3’ end of tracrRNA. In tr_L_^*^, tr_S_ transcription is below the limit of detection and tr_P_ levels are greatly reduced, likely owing to poor processing of tr_L_ compared to tr_S_.

**Figure S3.**
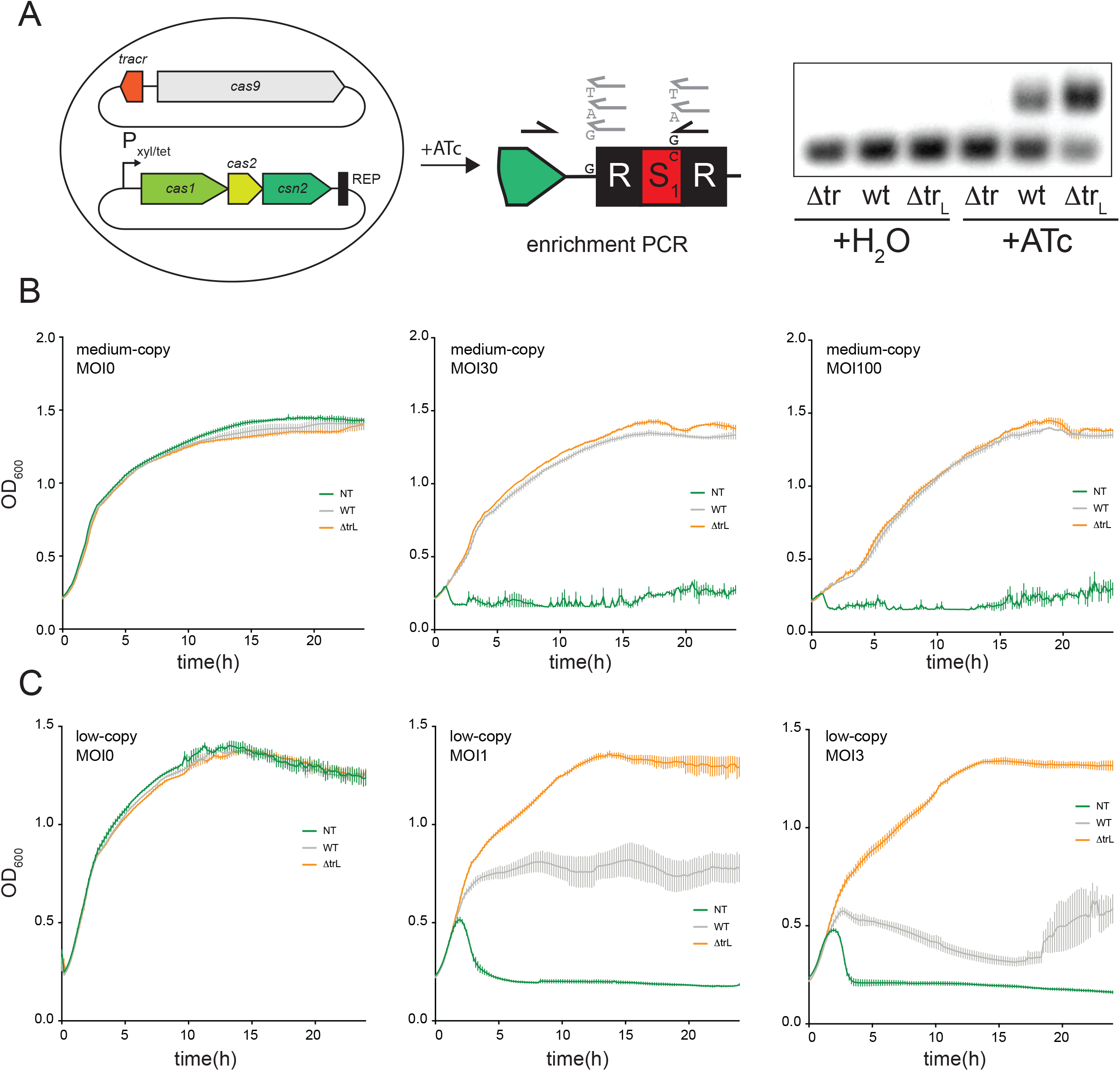
Δtr_L_ cells exhibit enhanced CRISPR adaptation and interference. **A)** Schematic of spacer acquisition assay on cells harboring a plasmid expressing tracrRNA and *cas9* and a second plasmid expressing the adaptation-specific Cas genes and CRISPR repeat from the anhydrotetracycline(ATc)-inducible promoter P_xyl/tet_. ATc was added at 0.5 ug/mL for 2 hours to induce expression of the adaptation cassette, and spacer acquisition was monitored by an “enrichment PCR” assay. PCR products were run on a 2% agarose gel, and the presence of a larger band represents a single newly acquired spacer. **B-C)** Interference assays on cells harboring CRISPR systems with the ϕNM4γ4-targeting spacer NM2 on medium (B) and low-copy (C) plasmids. Cells were treated with ϕNM4γ4 at the indicated MOIs and cell densities (OD_600_) were measured every 10 minutes in a 96-well plate reader.

**Figure S4.**
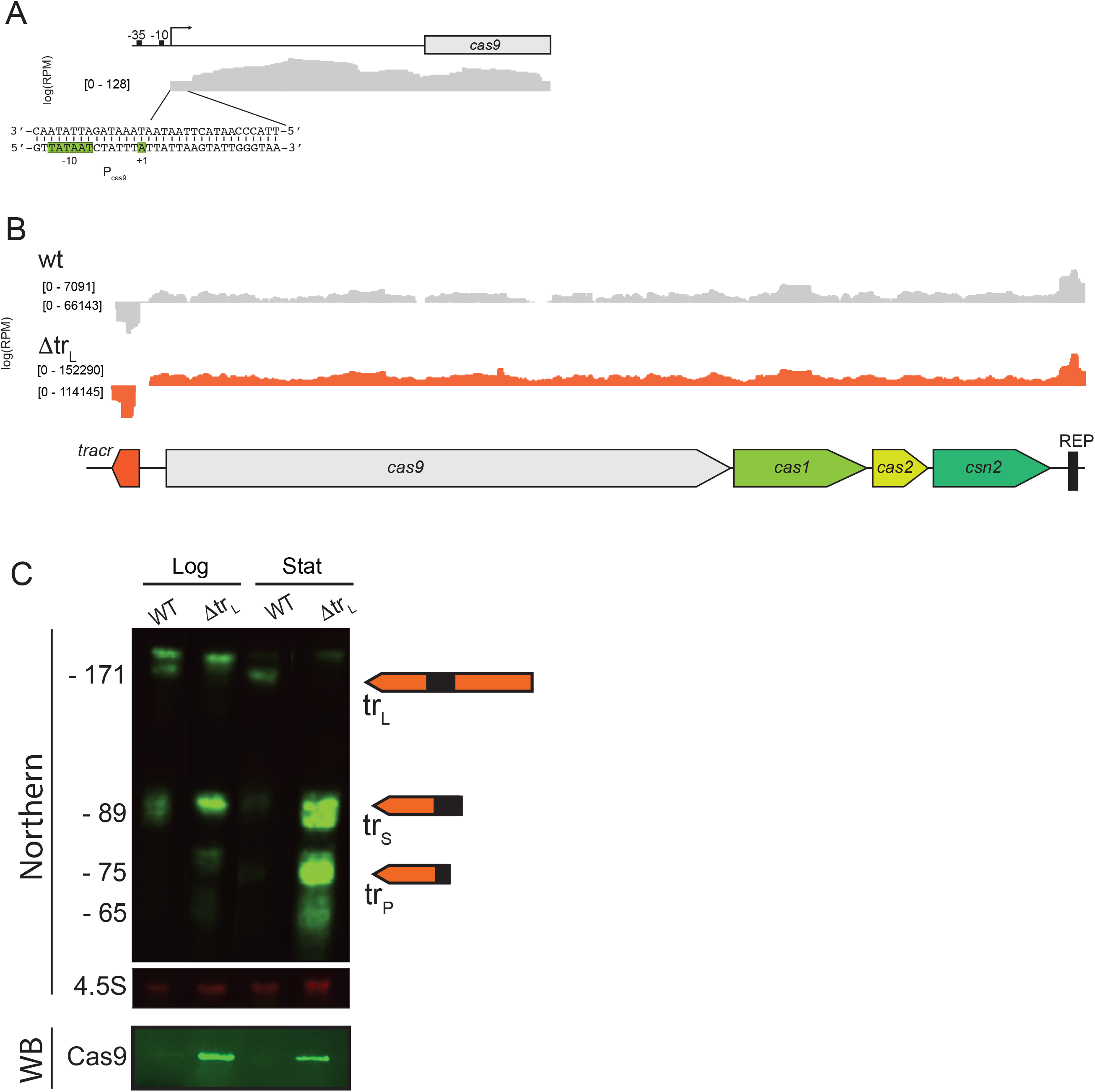
Transcriptomic changes in Δtr_L_. **A)** RNAseq data from cells harboring a Δtr_L_ CRISPR system outgrown from stationary to mid-logarithmic phase for 4 hours. A window is shown focused on the P_cas9_ region. **B)** RNAseq from cells with a wild-type or Δtr_L_ CRISPR system grown to mid-log phase as in (A) showing sequencing reads across the broader CRISPR-Cas region. Note the y-axis is scaled differently in the wild-type and Δtr_L_ experiments. **C)** Infrared Northern blot on strains from (B) grown overnight to late stationary phase (“stat”) or diluted back from stationary into logarithmic phase for 4 hours (“log”). Membranes were probed with oligos matching the 3’ end of tracrRNA (top panel) or the 4.5S RNA (middle panel). The bottom panel shows a Western blot (WB) for Cas9.

**Figure S5.**
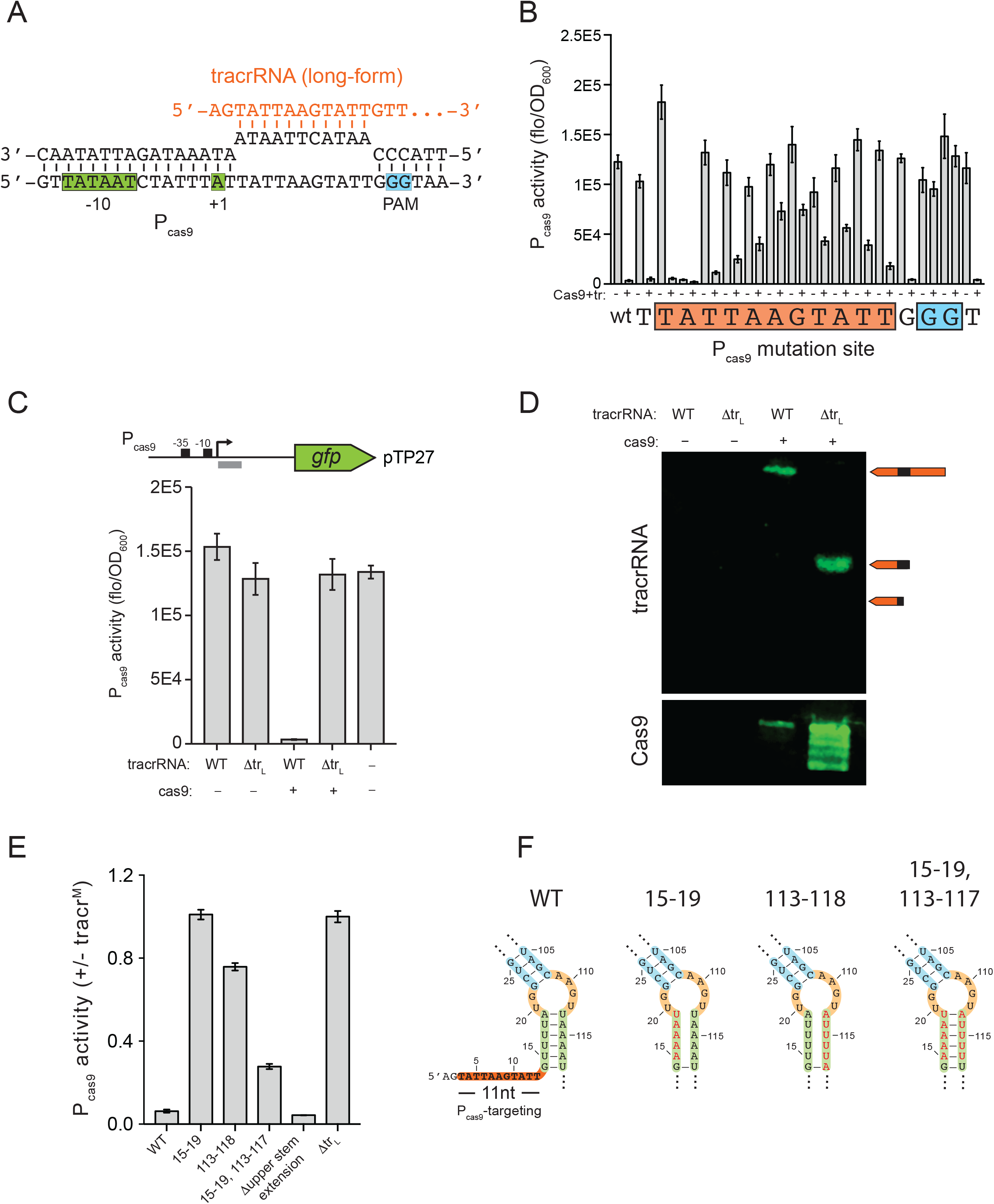
Extended characterization of the determinants of tr_L_:P_cas9_ repression. **A)** Base-pairing between the 5’ end of tr_L_ and the P_cas9_ region just downstream from the transcriptional start site (+1). −10 and +1 promoter elements are shown in green, and the PAM downstream from the tr_L_ targeted site is shown in cyan. **B)** P_cas9_ promoter activity was measured as in Fig. 4B with the absolute values for both +/-Cas9+tracrRNA(tr) conditions shown. **C)** Promoter activity was measured (fluorescence/OD_600_) in cells harboring a P_cas9_-GFP reporter plasmid and a second plasmid expressing wild-type tracrRNA or Δtr_L_ and wild-type cas9 or a cas9 null mutant in which two stop codons were inserted after the 15^th^ codon. **D)** Infrared Northern blot on the strains in (C) grown to stationary phase. Membranes were probed with oligos matching the 3’ end of tracrRNA. **E)** Promoter activity was measured (fluorescence/OD_600_) in cells harboring a plasmid expressing cas9 from a constitutive promoter and P_cas9_-GFP and a second plasmid expressing the indicated tracrRNA mutants. The ratio of activities for the indicated tracrRNA mutants relative to Δtr_L_ are shown. Numbers indicate the positions of nucleotides mutated to their complementary base relative to the 5’ end of tr_L_; Δupper stem extension; nucleotides 26 – 104 of tr_L_ were replaced by a 5’-GAAA-3’ tetraloop. **F)** Schematic showing the base-pairing potential of the tr_L_ mutants from (E).

**Figure S6.**
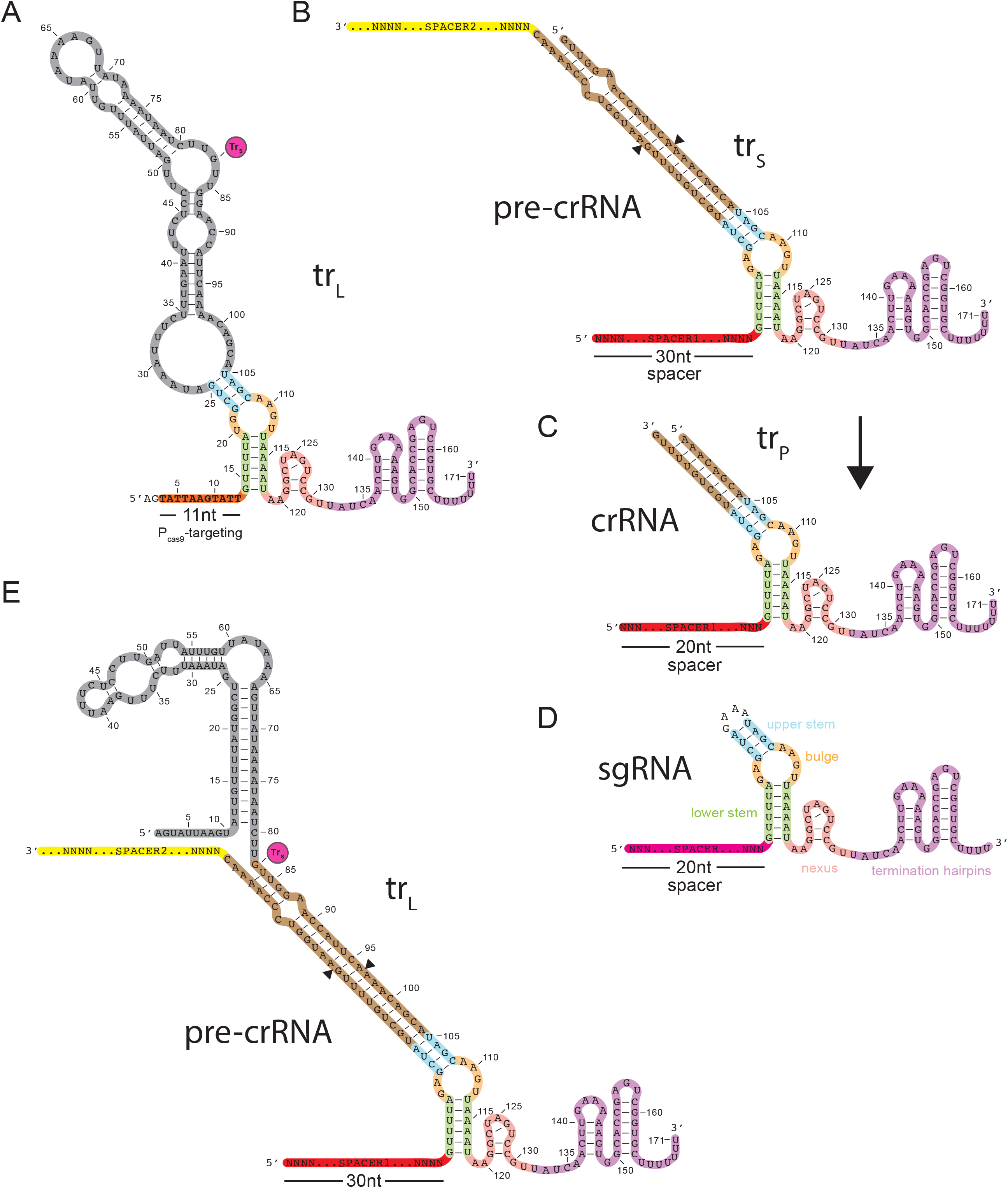
Structures of CRISPR-Cas RNAs. Schematics are shown for the putative tr_L_ natural sgRNA **(A)**, the tr_S_:pre-crRNA duplex **(B)**, the processed tr_P_:crRNA duplex **(C)**, the artificial sgRNA commonly used in CRISPR editing technologies **(D)**, and the tr_L_:pre-crRNA duplex **(E)**. Green, lower stem; dark yellow, bulge; cyan, upper stem; pink, nexus; purple, termination hairpins; brown, region of crRNA:tracrRNA complementarity outside the stems; gray, putative structures of tr_L_ regions of unknown function.

**Figure S7.**
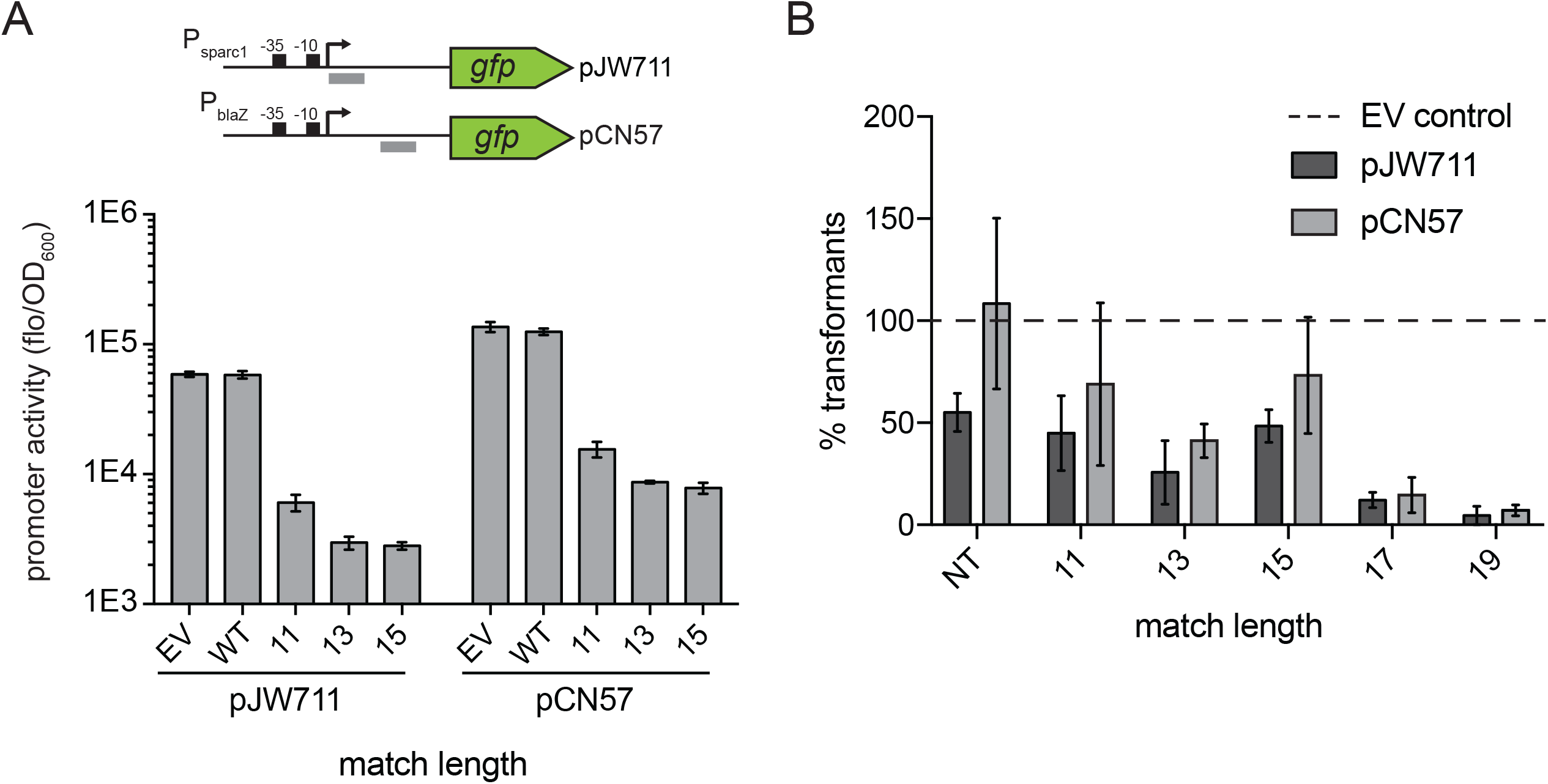
Extended tr_L_ match lengths lead to tighter repression and cleavage. **A)** Promoter activity was measured (fluorescence/OD_600_) in cells harboring variants of pRW22 (Fig. 5A) with target site matches of the indicated lengths and a second GFP reporter plasmid, pJW711 or pCN57. Grey bars in the schematic, targeting site; EV, empty vector; WT, wild-type tr_L_ targeting P_cas9_. **B)** Cells harboring pRW22 variants with the indicated target site match lengths were transformed with a second plasmid, EV, pJW711 or pCN57. Transformants were plated on selective media with antibiotics to maintain both plasmids, and surviving colonies were counted. Data is presented as a ratio of pJW711 or pCN57 colonies relative to the EV control.

**Figure S8.**
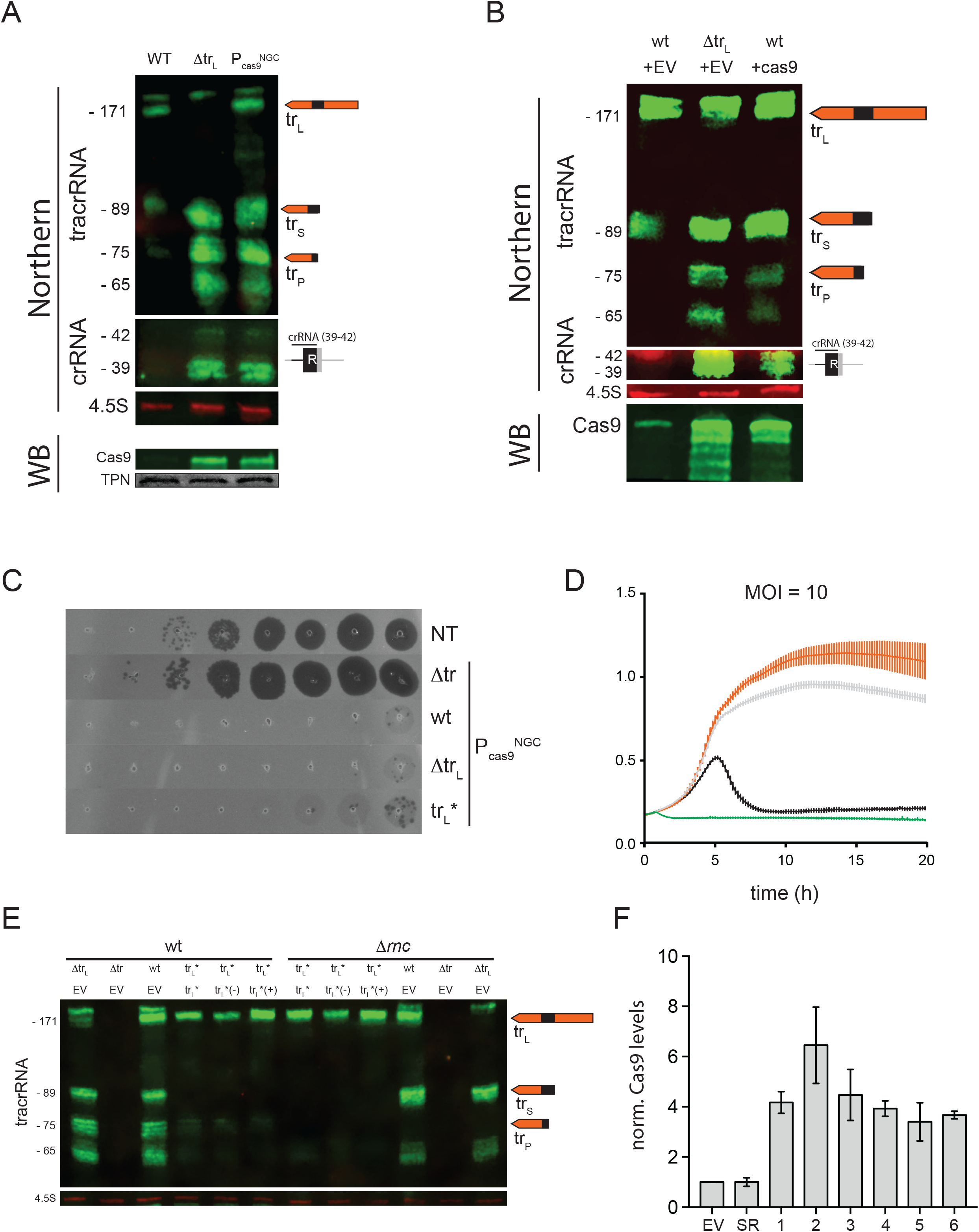
crRNAs interact with tr_L_ to regulate CRISPR-Cas levels and activity. **A-B)** Infrared Northern blot of (A) strains from Fig. 6A or (B) cells harboring a plasmid with a naïve wild-type or Δtr_L_ CRISPR system and a second empty vector or a plasmid constitutively expressing cas9 from P_spac_ (+cas9). Membranes were probed with oligos matching the 3’ end of tracrRNA (top), the crRNA repeat (middle) or the 4.5S RNA (bottom, loading control). Bottom panels, Western blot (WB) using an α-Cas9 antibody. Membranes were stained with Ponceau S, and the prominent band is shown as a loading control (total protein, TPN). **C)** Top agar interference assay with cells harboring a plasmid expressing Cas9, the indicated tracrRNA mutants and the ϕNM4γ4-targeting spacer NM2. 10-fold dilutions of ϕNM4γ4 were plated on the indicated bacterial lawns. Δtr, deletion of the tracrRNA locus; wt, wild-type tracrRNA; Δtr_L_, deletion of the long-form tracrRNA; tr_L_*, P_trS_ mutant expressing long-form tracrRNA only; P_cas9_^NGC^, P_cas9_ PAM mutant that is insensitive to tr_L_ repression. **D)** Interference assay with ϕNM4γ4 at MOI = 10 with the indicated strains from Figure 6B. IPTG was added at 1 mM to induce expression from P_sparc2_. **E)** Infrared Northern blot on cells harboring a plasmid expressing cas9 and the indicated tracrRNA mutants from the P_cas9_^NGC^ promoter and a second plasmid expressing additional tr_L_ or an empty vector. Membranes were probed with oligos matching the 3’ end of tracrRNA. tr_L_*(-)/tr_L_*(+), cells harboring plasmids expressing tr_L_* from the P_sparc2_ IPTG-inducible promoter were treated without or with IPTG for 1:15 hours respectively. **F)** Quantification of Western blot experiments described in Figure 6C, performed in biological triplicate.

**Figure S9.**
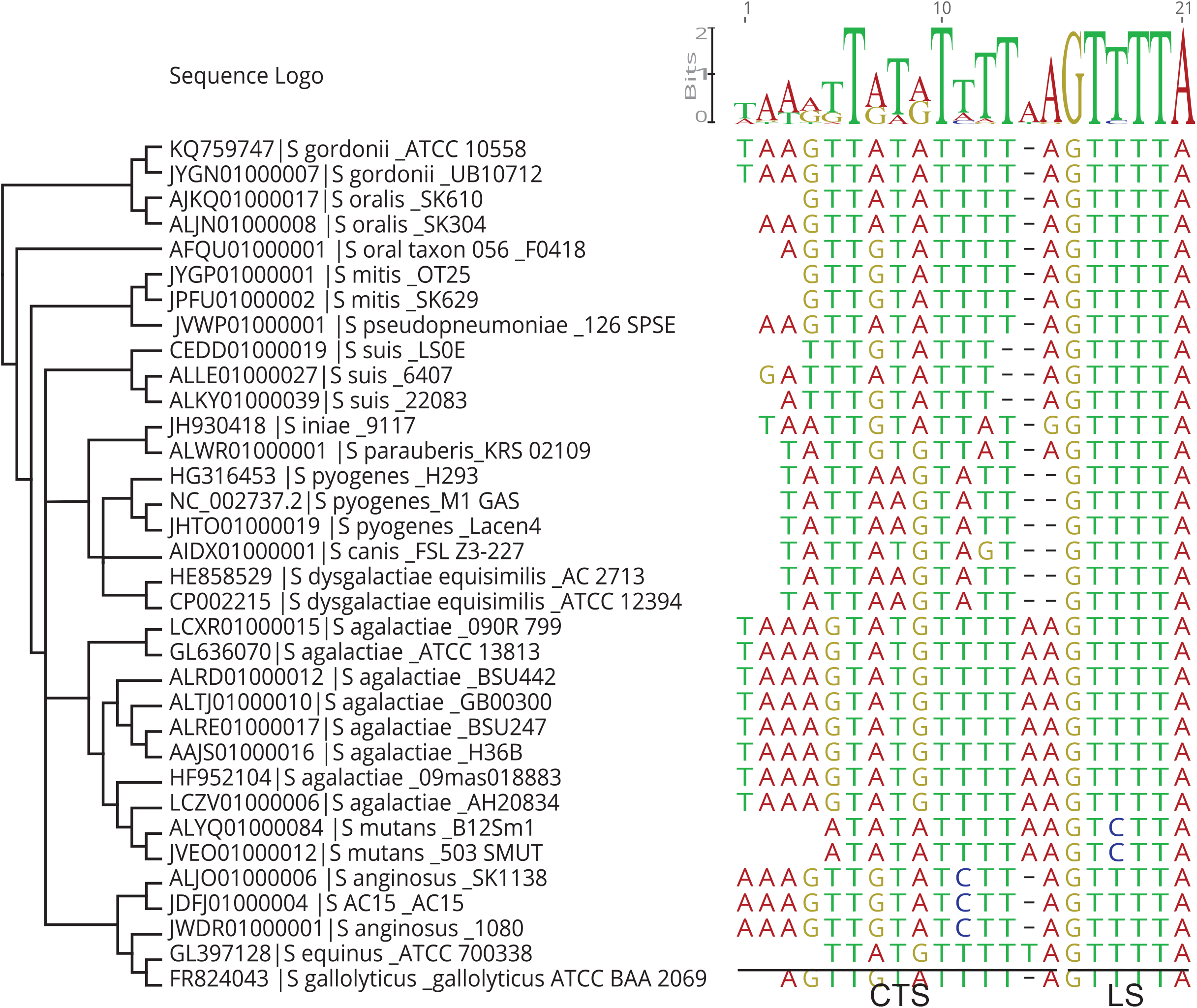
Sequence alignment of tr_L_ targeting determinants. A MAFFT multiple sequence alignment was constructed of the candidate targeting sequences (CTS) and lower stems (LS) from all tr_L_^+^ genomes. The phylogenetic tree was constructed as for Figure 7 using Cas9 protein sequences.

**Figure S10.**
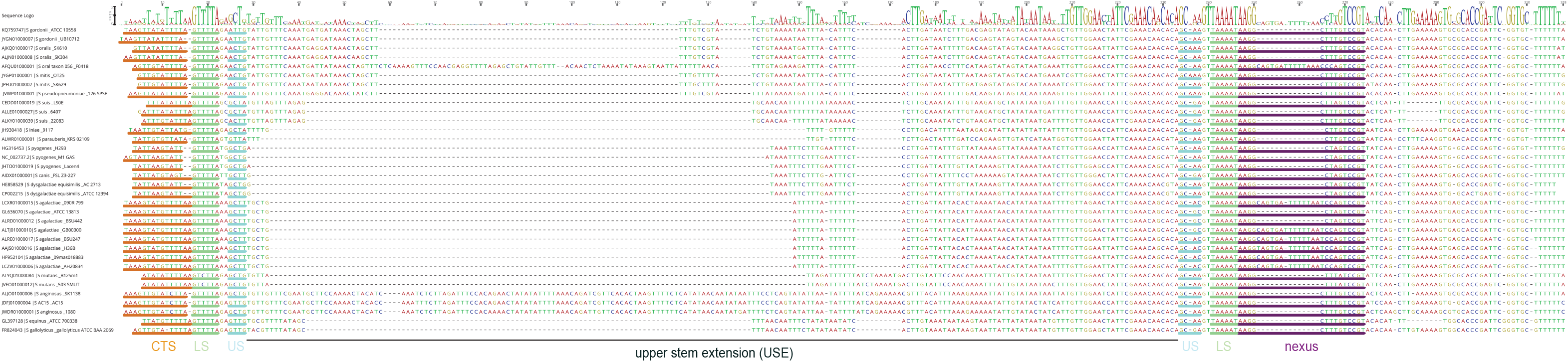
Sequence alignment of full-length tr_L_. A MAFFT multiple sequence alignment was constructed of tr_L_ from all tr_L_^+^ genomes. Orange, candidate targeting sequence (CTS); green, lower stem (LS); blue, upper stem (US); black line, upper stem extension (USE); purple, nexus.

**Figure S11.**
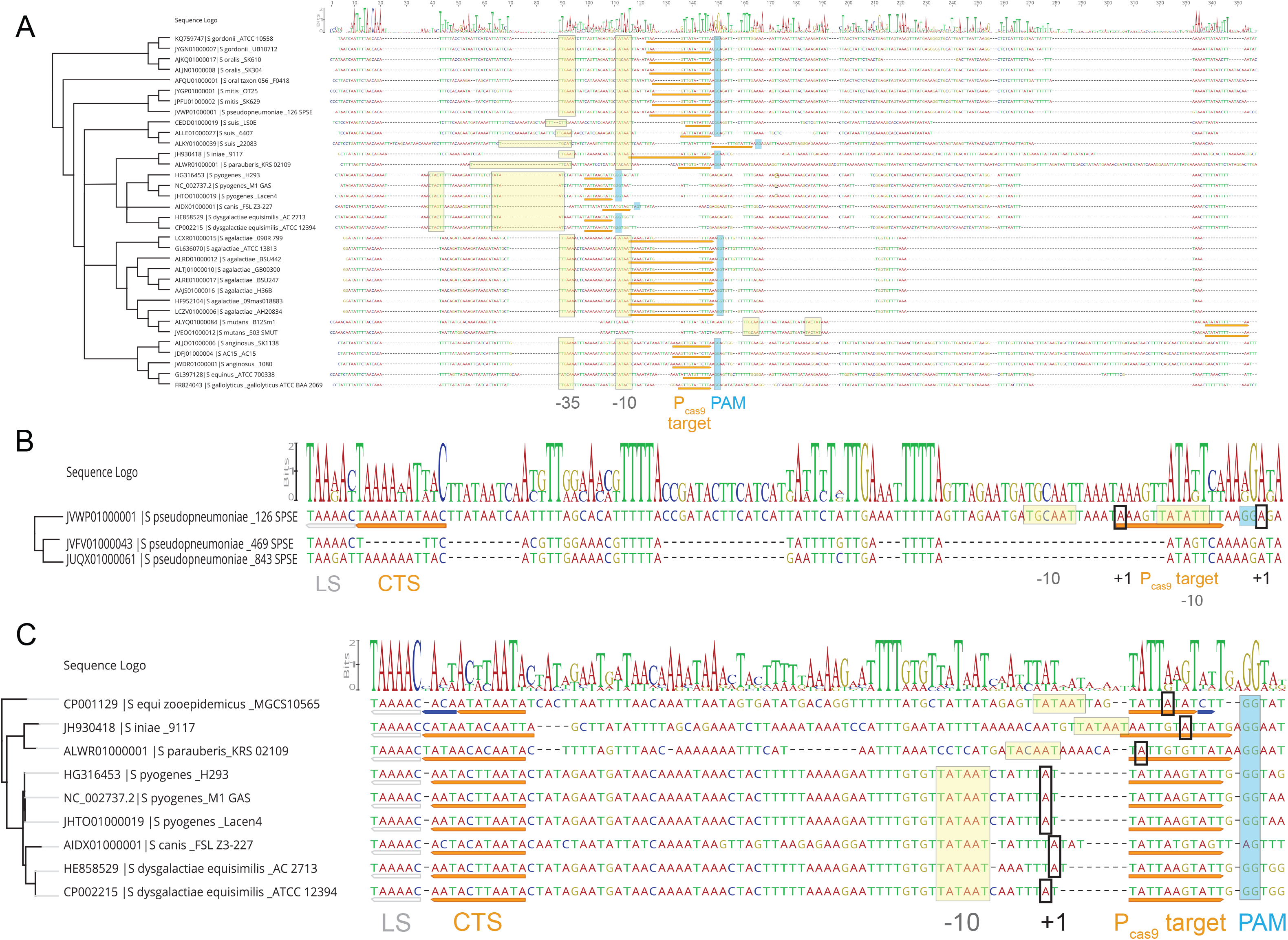
Evidence of variation and degeneration within Pcas9 and its targeting determinants. **A)** A MAFFT multiple sequence alignment was constructed for the P_cas9_ regions of tr_L_^+^ genomes. Yellow boxes, predicted −35 and −10 P_cas9_ promoter elements; orange underline, P_cas9_ target site; light blue boxes, PAM. **B-C)** A multiple sequence alignment spanning the tr_L_ candidate targeting sequence (CTS, orange underline) and lower stem (LS, grey underline) through the P_cas9_ target site in several *S. pseudopneumoniae* strains (B) and a *S. pyogenes* local branch (C). Yellow and black boxes, putative −10 and +1 promoter elements respectively. In *S. equi zooepidemicus*, the targeting determinants contain seed mismatches (blue underline) but retain an LS and PAM.

**Supplemental Table 1.** An Excel file containing NGS data from the experiments detailed in Figure 1. Separate worksheets are provided showing NGS reads that mapped to the genome and CRISPR plasmid separately. The - CRISPR library was treated without phage in single and with phage in duplicate. The +CRISPR library was treated without phage in single and with phage in triplicate.

**Supplemental Table 2.** An Excel file containing metadata for the bioinformatic analysis described in Figure 7.

**Supplemental Table 3.** An Excel file containing RNA-seq data for the experiments described in Fig. S4.

**Supplemental Materials.** An Excel file with three worksheets detailing the strains, oligos and plasmids used in this paper. Plasmid construction details are also provided.

## METHODS

### Growth conditions

*Staphylococcus aureus* strains were grown at 37°C, unless otherwise indicated, in Bacto Brain-Heart infusion (BHI) broth with shaking at 220 RPM. During outgrowths from stationary phase preceding phage treatments, BHI was supplemented with calcium chloride at 5 mM to allow phage adsorption and with 1 mM IPTG to allow expression from P_sparc2_ when necessary. Antibiotics were used at the following concentrations for strain construction and plasmid maintenance in *S. aureus*: tetracycline, 5 μg/mL; chloramphenicol, 10 μg/mL; erythromycin, 10 μg/mL; spectinomycin, 250 μg/mL.

### Strain and plasmid construction

See Supplemental Materials for strains, plasmids, cloning notes and oligos used in this study.

### Gibson assembly

Gibson assemblies were performed as described^1^. Briefly, 100 ng of the largest dsDNA fragment to be assembled was combined with equimolar volumes of the smaller fragment(s) and brought to 5 μL total in dH_2_O on ice. Samples were added to 15 μL of Gibson Assembly master mix, mixed by pipetting and incubated at 50°C for 1 hour. Samples were drop dialyzed in dH_2_O for 30 minutes – 1 hour, and 5 μL were electroporated into 50 μL electrocompetent RN4220 *S. aureus* cells.

### Oligo cloning

To create a repeat-spacer-repeat CRISPR array with a defined spacer, we used a restriction digest-based cloning approach. Parent plasmids contain two CRISPR repeats flanking a 30bp sequence housing two BsaI restriction sites. 400-800 ng of the plasmid was mixed with the BsaI-HFv2 restriction enzyme (NEB, R3733S) in a 10 μL reaction volume (1 μL BsaI-HFv2 enzyme, 1 μL CutSmart buffer, plasmid + nuclease-free water to 10 μL) and incubated at 37°C for ~8 hours. Two IDT oligos, a “top” strand with sequence 5’-AAAC-(30bp spacer)-G-3’, and a “bottom” strand with sequence 5’-AAAAC-(30bp spacer reverse complement)-3’ were phosphorylated with PNK (NEB, M0201S) in a 50 μL reaction volume (1.5 μL 100 uM top oligo, 1.5 μL 100 uM bottom oligo, 41 μL nuclease free water, 5 μL T4 PNK 5x reaction buffer, 1 μL T4 PNK) at 37°C for 30 minutes – 1hr. After phosphorylation, oligos were annealed by adding 2.5 μL of 1M NaCl to the 50 μL reaction and incubating for 5 minutes at 98°C, then allowing the reaction to cool to room temperature (1-2 hours). The phosphorylated, annealed oligos were diluted 1:10 in nuclease-free water and ligated to the digested plasmid in a 20 μL reaction (10 μL digested plasmid, 6 μL water, 1 μL 1:10 diluted oligos, 2 μL T4 ligase buffer, 1 μL T4 DNA ligase enzyme (NEB, M0202S)) at room temperature overnight. Reactions were drop dialyzed for 1 hour in dH_2_O and 5 μL were transformed into electrocompetent RN4220 *S. aureus* cells.

### Tn-seq screen

To construct a transposon library in cells lacking a CRISPR system, *Staphylococcus aureus* RN4220 cells harboring pTV1, a temperature-sensitive plasmid containing transposon Tn917 from *Streptococcus faecalis^2^*, were grown overnight at the permissive temperature (30°C) in BHI supplemented with chloramphenicol. The next morning, 500 μL cells were washed 1x in plain BHI and then diluted into 500 mL BHI prewarmed to the restrictive temperature (42°C) and supplemented with erythromycin, without chloramphenicol. Cells were grown at 42°C with shaking for ~8 hours and diluted again 500 μL into 500 mL BHI/erythromycin. After an overnight growth at 42°C, cells were plated onto BHI agar plates supplemented with erythromycin and incubated overnight at 42°C. The next day, 10,000 colonies were scraped with 3 mL BHI into a Falcon tube and resuspended by pipetting and vortexing. DMSO was added to 10% and library aliquots were stored at −80°C. A second transposon library was constructed in cells harboring the CRISPR-Cas system on plasmid pJW92 as above, with tetracycline added during each step for plasmid maintenance.

Transposon library aliquots were thawed and diluted to OD = 0.05 in 20 mL (-phage experiments) or 500 mL (+ phage experiments) BHI supplemented with calcium chloride. After ~ 2 hours of growth at 37°C, culture ODs were roughly 0.4-0.5, and ϕNM4γ4 was added to the + phage experiments in duplicate (CRISPR-library) or triplicate (CRISPR+ library) at MOI = 1. After 24 hours of growth, genomic DNA was prepared using the Wizard Genomic DNA purification kit (Promega) and culture aliquots were frozen at −80°C for storage in 10% DMSO. As above, tetracycline was added in all steps for cells harboring pJW92. To prepare NGS libraries, genomic DNA was digested with NEBNext dsDNA Fragmentase (M0348) for 9 minutes, and the reaction was stopped by addition of 0.1 M EDTA (final concentration). The fragmented DNA, centered at 700bp, was then purified using the QIAquick PCR Purification kit and end repaired using the NEBNext DNA Library Reagent Set for Illumina (E6000S). After another QIAquick PCR purification, NEBNext adapters were ligated onto the fragments and DNA in the 0.2 – 1 kb range was purified from an agarose gel with the QIAquick Gel Extraction kit. Next, for each sample, a PCR was performed with a unique oligo from the NEBNext Multiplex Oligos for Illumina kit (E7335S, Index Primers Set 1) and a universal, transposon-specific primer (oJW451). Amplicons in the 0.2 – 0.5 kb range were gel purified as before and sequenced on an Illumina HiSeq using the sequencing primer oJW436. Sequencing reads were processed with custom Python scripts and aligned to the NCTC8325 genome using the Burrows-Wheeler Aligner. Tn-seq output was visualized with the Integrative Genomics Viewer^3^.

### *S. aureus* miniprep protocol

1-1.5 mL of an overnight culture, unless otherwise indicated, was pelleted and resuspended in 250 μL Buffer P1. 10-20 μL lysostaphin (Ambi Products LLC, LSPN-50, 100 μg/mL final) was added and the cells were incubated without shaking at 37°C for ~20 minutes. Following lysis, plasmids were isolated using the QIAGEN Spin Miniprep kit according to the manufacturer’s protocol, beginning with addition of P2. DNA was eluted from each column in 30 μL RNAse-free water.

### Electroporation adaptation assay

Electrocompetent *S. aureus* cells were made by washing overnight cultures twice in full-volumes of dH_2_O and once in a half-volume of dH_2_O before resuspending in 1/100 the original volume in 10% glycerol. 50 ng of dialyzed DNA, amplified from phage ϕNM4γ4 with primers JW1744 and JW1745, were added to 50 μL of electrocompetent cells in an Eppendorf tube, mixed by pipetting and left at RT for 10 minutes. Cells were transferred to a 0.2 cm electroporation cuvette (Bio-Rad, 1652086) and electroporated at 1.8 kV using an Eporator (Eppendorf). Following electroporation, 1 mL BHI broth was added and the contents of the cuvette were mixed by pipetting and then transferred to an Eppendorf tube and grown for 1 hour with shaking at 220 RPM at 37°C. Plasmids were miniprepped, and spacer acquisition was monitored by a spacer-specific PCR reaction with primers oJW1833 and oJW1834. PCR products were visualized on a 2% agarose gel run for 25 minutes at 130 V.

### Overexpression adaptation assay

Overnight cultures of *S. aureus* were diluted to OD = 0.04 in BHI broth supplemented with antibiotics and grown for 1 hour with shaking at 37°C. Cultures were then supplemented with anhydrotetracycline (ATc) at 0 μg/mL or 0.5 μg/mL (final concentrations). Cultures were incubated with shaking at 37°C for 2 hours and then miniprepped with a lysis step including 15 μL of lysostaphin incubated at 37°C for 5 minutes. An enrichment PCR was performed as described^4,5^ and PCR products were visualized on a 2% agarose gel run for 27 minutes at 130 V.

### Liquid growth interference assay

Overnight cultures of *S. aureus* were diluted to OD = 0.025 in BHI broth supplemented with calcium chloride and antibiotics and grown for 1 hour and 15 minutes shaking at 37°C. Cultures were normalized to the OD of the lowest culture and phage ϕNM4γ4 was added to the appropriate MOI. After inverting to mix, 150 μL of each culture were added to a flat-bottom 96-well plate (Grenier 655180), and the plate was incubated at 37°C with shaking in a TECAN Infinite F Nano+ with OD_600_ measurements recorded every 10 minutes for 24 hours.

### Top agar interference assay

100 μL of *S. aureus* overnight cultures were added to a 50 mL Falcon tube. 6 mL of BHI top agar (0.75% agar) supplemented with calcium chloride (5mM final concentration) were added to each tube. After swirling to mix, the cells and top agar were poured onto a BHI 1.5% agar plate and rocked gently to create a bacterial lawn. The plate was incubated for 15 - 30 minutes at RT. 3.5 μL of 8 10-fold serial dilutions of phage ϕNM4γ4 in BHI broth were spotted on top of the bacterial lawn using a multichannel pipette. After a 30 minute incubation at RT, the plates were moved to a 37°C incubator overnight.

### Promoter activity fluorescence assay

200 μL of each overnight culture was spun down in a 1.5 mL Eppendorf tube at 6,000 RPM for one minute. Cell pellets were resuspended in 1 mL of 1X PBS and 150 μL were transferred into a clear, flat-bottomed 96-well plate (Grenier 655180). Measurements for absorbance (at 600 nm) and fluorescence (excitation wavelength = 485 nm; emission wavelength = 535 nm) were taken using a TECAN Infinite F Nano+. For each experimental strain, promoter activity was measured as (Fe)/(Ae) - (Fc)/(Ac) where F = fluorescence, A = absorbance, e = experimental strain and c = non-fluorescent control strain.

### Liquid growth immunity assay

Overnight cultures of *Staphylococcus aureus* harboring naïve CRISPR systems with a single repeat were diluted to OD = 0.1 in BHI supplemented with calcium chloride and relevant antibiotics and grown for 1.5 hours. Cultures were normalized to the OD of the lowest culture, and phage ϕNM4γ4 was added at the specified MOIs. 150 μL of each culture was added to a clear, flat-bottom 96-well plate (Grenier 655180), and the plate was incubated at 37°C with shaking in a TECAN Infinite F Nano+. OD_600_ measurements were recorded every 10 minutes for 24 hours.

### Top agar immunity assay

100 μL of overnight cultures of *Staphylococcus aureus* harboring naïve CRISPR systems were mixed with ϕNM4γ4 at MOI = 25 and 6 mL of BHI top agar (0.75% agar) supplemented with calcium chloride and poured onto BHI agar (1.5%) plates. After 15 minutes at RT, plates were moved to a 37°C incubator overnight, and surviving colonies were counted. When indicated, colony PCR was performed with primers oJW154 and oJW355 to verify expansion of the CRISPR array due to spacer integration.

### PCR conditions

PCR was performed with Phusion HF DNA polymerase using 5X Phusion Green Reaction Buffer (Thermo). Each reaction contained 10 μL buffer, 4 μL dNTPs, 0.5 μL each of 100 μM forward and reverse primers, 10-50 ng template, 0.5 μL polymerase and nuclease-free water to 50 μL. Three-step cycling was performed under the following conditions: 98°C for 30 seconds, 34 cycles of [98°C 5 seconds, 45-72°C 15 seconds, 72°C for 30s/kb], 72°C 10 minutes, hold at 10°C.

### RNA extraction

To extract RNA for Northern blot, RT-qPCR, and RNA-seq analysis, 7.5E8 cells from an overnight culture were spun down, resuspended in 150 μL 1X PBS (10X stock, Corning, 46-013-CM) and lysostaphin (60 μg/ml final concentration), and incubated at 37°C for 5 minutes. To the whole cell lysate, 450 μL Trizol (Zymo, R2071) and 600 μL 200 proof ethanol were added, samples were vortexed and RNA was extracted using the Direct-Zol Miniprep Plus spin column according to the manufacturer’s protocol (Zymo, R2071). Samples were eluted in 50 μL Ambion RNAse-free water (ThermoFisher, AM9937).

### Infrared Northern (irNorthern)

Total RNA (3-10 μg) was mixed 1:1 with 2X Novex sample buffer (Thermo, LC6876), boiled at 94°C for 3 minutes and placed on ice for 3-5 minutes. Samples were loaded onto a 15% TBE-Urea gel (MINI-Protean, Bio-rad, 4566053) and run at 150 V for 2-2.5 hours. A Hybond N+ membrane (GE lifesciences, 45000854) and 6 sheets of 3 mm Whatman cellulose paper (Sigma Aldrich, WHA3030861) were pre-soaked for 5 minutes in room temperature 0.5X TBE, then assembled into a sandwich: 3 layers Whatman paper, Hybond membrane, TBE-Urea gel, and 3 more layers of Whatman paper. Blotting was performed using a Trans-blot Turbo (Bio-Rad) at 200 mA for 30 minutes. The membrane was then pre-hybridized in 10 mL ExpressHyb (Clontech, NC9747391) at 44°C in a rotating oven for 1 hour, and probed overnight at 44°C, with rotation, using probes conjugated to irDyes (LICOR; 680CW 929-50010, 800CW 929-50000), made following the protocol detailed at (https://bio-protocol.org/e3219)^6^. The membrane was washed once with 2X SSC/0.1% SDS, and once with 1X SSC/0.1% SDS each for 10 minutes at RT, then visualized on the Odyssey Fc (LICOR). 4.5S RNA was used as a loading control (stability of 4.5S across genetic backgrounds was verified by qPCR). Sequences for oligos used to probe crRNA, tracrRNA and 4.5S RNA (oJW2313, oJW1991, oJW2172) are listed in Supplementary Materials.

### Western blot

1.8E8 cells were resuspended in 1X PBS supplemented with lysostaphin (60 μg/ml final concentration) and incubated at 37°C for 20 minutes. Whole cell lysate was mixed 1:1 with 2x Laemmli solution (Bio-rad, 1610737) supplemented with B-mercaptoethanol (55 mM stock, ThermoFisher, 21985023) at a final concentration of 1.3 mM, and boiled at 98°C for 10 minutes. Samples were loaded onto a 4-20% Tris-glycine gel (MINI-Protean TGX Pre-cast, Bio-rad, 4561095) and run at 200 V for 15 min-1 hour. A PVDF membrane (Bio-rad) was hydrated with methanol for 15-30 seconds and pre-wet alongside a stack of blotting paper for 3-5 minutes in 1X Transfer buffer. A blotting sandwich was assembled consisting of six layers (one stack) of filter paper, the PVDF membrane, Tris-glycine gel, and another stack of filter paper. Samples were transferred onto a nitrocellulose membrane with the Trans-blot turbo (Bio-Rad, 1704150), with high molecular weight transfer settings (1.6 A for 10 minutes), and the membrane was stained with Ponceau for 5 minutes to perform total protein normalization. The membrane was blocked with 5% nonfat dry milk for 1 hour, then probed with a 1:1000 dilution of Cas9 monoclonal antibody (Cell Signaling, 7A9-3A3) for 2 hours at RT, or at 4°C overnight. The membrane was washed with 1X TBST buffer 3x for 10 minutes at RT, then probed with 1:15,000 dilution of infrared (LICOR, 925-32210) or 1:10,000 dilution of HRP (Pierce, PA174421) secondary antibody for 1 hour at room temperature. The membrane was washed as before, then visualized on the Odyssey Fc (LICOR).

### RNA-seq library preparation and sequencing

To prepare libraries for next-generation sequencing, we first depleted ribosomal RNA from the total purified RNA using reverse capture of biotinylated probes annealing to rRNA with streptavidin beads, following the protocol detailed at (https://mbio.asm.org/content/11/2/e00010-20)^7^. rRNA depleted samples were prepared for sequencing using the Cleantag Small RNA library prep kit (Trilink, L-3206-24) according to the manufacturer’s instructions, with 10 ng RNA input and 18 PCR cycles. Sequencing was performed on an Illumina MiSeq with a v3 2×75 cycle kit at the Johns Hopkins Genome Resources Core Facility (GRCF). Full data is provided in Supplemental Table 3.

### qPCR

Purified total RNA (1-4 μg) was treated with TURBO DNA-free (ThermoFisher, AM1907) and reverse transcribed with Superscript IV (ThermoFisher, 18090050) according to the manufacturer’s instructions. cDNA was diluted to 0.5-1 ng/μL and 4-8 ng was used as input in an 8 μL volume. 1 μL of 10 μM forward/reverse primers were used, and 10 μL of 2X PowerUp SYBR mastermix (ThermoFisher, A25742). qPCR was performed with cycling conditions: 50°C for 2 minutes, 95°C for 2 minutes, 39 cycles of [60°C 1 minute], followed by a melt curve: 65°C to 95°C, incrementing 0.5°C every 5 seconds. *rho* was used as a loading control and was amplified using primers oJW2003/2004. *cas9* was amplified with primers oW262/oJW1986. *cas1* was amplified using primers oJW150/oL432. *csn2* was amplified using primers oJW2139/2140. The region spanning *csn2* and *crRNA* (*csn2_cr*) was amplified using oJW2141/2142. The *pre-crRNA* was amplified using oJW2142/2143. All forms of *tracrRNA* were amplified using oJW2012/2013. qPCR primer sequences are provided in Supplementary Materials.

### Transformation assay

10 mL overnight cultures of cells harboring derivatives of pTP16, which encodes *cas9* and *tracrRNA* variants with increasing match lengths to the GFP reporter plasmids pJW711 and pCN57 were diluted to OD = 0.1 and outgrown in BHI supplemented with chloramphenicol for 1-2 hours until the OD was between 0.8-1. The cultures were centrifuged at 4200 RPM for 10 minutes and washed twice with 1 mL of ice-cold deionized water in a 1.5 mL Eppendorf tube with 6000 RPM 1 minute spins between washes. Cells were resuspended in 150 μL 10% glycerol and stored at −80°C if not used immediately. 50 ng of pJW711, pCN57, or an empty vector pE194 was electroporated into 50 μL of competent cells, and outgrown in 300 μL BHI at 37° C with shaking for 3 hours. 100 μL of each outgrowth was plated onto BHI agar plates supplemented with chloramphenicol and erythromycin and total transformants were counted the following day.

### *In vitro* transcription (IVT)

IVT templates were generated using the templates pTP16 and pRW22-26 with forward primers oJW2268-2274, each of which begin with the T7 promoter sequence (5’-GAAATTAATACGACTCACTATAGG-3’), and the reverse primer oJW2267. To generate the WT long-form tracr, pTP16 was amplified with oJW2267 and oJW2268. To generate the GFP-targeting tracr with an 11-bp match, pRW22 was amplified with oJW2267 and oJW2269. To generate the GFP-targeting tracr with a 13-bp match, pRW23 was amplified with oJW2267 and oJW2270. To generate the GFP-targeting tracr with a 15-bp match, pRW24 was amplified with oJW2267 and oJW2271. To generate the GFP-targeting tracr with a 17-bp match, pRW25 was amplified with oJW2267 and oJW2272. To generate the GFP-targeting tracr with a 19-bp match, pRW26 was amplified with oJW2267 and oJW2273. To generate the sgRNA version of the GFP targeting tracr with a 20-bp match, pRW27 was amplified with oJW2267 and oJW2274. tracrRNA (wild-type and variants) was *in-vitro* transcribed from 1 μg of these templates using the HiScribe T7 High Yield RNA synthesis kit (NEB, E2040S). Reactions were incubated at 37°C for 4 hours and run on a 6% TBE-Urea gel (Invitrogen, EC6265BOX). Bands of the appropriate size were excised, added to a sterile Eppendorf tube in 450 μL gel extraction buffer (1X: 0.3 M NaOAc, 10 mM Tris-Cl pH 7.5, 1 mM EDTA), and incubated overnight at 4°C in an end-over-end rotator. The buffer with dissolved gel slice was added to 1 mL ice-cold 100% ethanol and 2 μL of glycogen (20 mg/ml, ThermoFisher, R0561) and incubated at −20°C for at least one hour. Samples were centrifuged at max speed at 4°C for 30 minutes, the supernatant was decanted, and the pellet was washed with 1 mL cold 70% ethanol. After another centrifugation step at max speed for 10 minutes at 4°C, the pellets were either dried in a vacuum centrifuge or let air dry for 10 minutes and eluted in Ambion RNAse-free water.

### Electrophoretic mobility shift assay (EMSA)

DsDNA targets were prepared by mixing 20 μL of 20 μM top and bottom strand oligos (Pcas: oJW2507/2508, Pgfp:oJW2509/2510) in annealing buffer (10 mM Tris pH 7.5, 50 mM NaCl, 1 mM EDTA), heating to 95°C for 5 minutes, then slowly cooling the mixture to room temperature for 1-2 hours. Annealed oligos were mixed with 4 μL 6X loading dye (NEB, B7024S), loaded onto a lab-made 8% TBE gel (Acrylamide/Bis solution 37.5:1, Bio-rad, 1610148), run at 200V for 1 hour, then stained with SYBR Gold (ThermoFisher, S11494) for 10 minutes still, and 10 minutes shaking. Bands of the appropriate size were excised and added to a sterile Eppendorf tube in 450 μL gel extraction buffer (1X: 0.3 M NaOAc, 10 mM Tris-Cl pH 7.5, 1 mM EDTA), and incubated overnight at room temperature in a shaker. Gel extraction buffer was added to 1 mL ice-cold 100% ethanol and 2 μL of glycogen (20 mg/mL, ThermoFisher) and incubated at −20°C for at least one hour. Samples were centrifuged at max speed at 4°C for 30 minutes, and the pellet was washed with 1 mL cold 70% ethanol. Following another centrifugation step at max speed for 10 minutes at 4°C, pellets were either dried in a vacuum centrifuge or let air dry for 10 minutes, then eluted in RNAse-free water (Ambion). Purified dsDNAs were diluted to 200 nM with binding buffer (20 mM HEPES pH 7.5, 250 mM KCl, 2 mM MgCl2, 0.01% Triton X-100, 0.1 mg/mL BSA, 10% glycerol), then diluted again 1:10 in binding buffer to make a 20 nM stock.

After dsDNA purification, the substrate was radiolabeled with P32 (1 μL 20 nM oligo duplex, 5 μL T4 PNK buffer, 10 mM radioactive ATP, 2 μL T4 PNK in a 50 μL reaction). This reaction was incubated at 37°C for 1 hour and cleaned up with ProbeQuant G50 spin columns (GE Healthcare, GE28-9034-08) as per manufacturer’s instructions.

Wild-type tr_L_ was prepared by performing *in vitro* transcription as described in *in vitro* transcription methods above. Tr_L_ was diluted to 8 μM in RNA hybridization buffer (20 mM Tris-HCl, pH 7.5, 100 mM KCl, 5 mM MgCl2), then heated at 95°C for 30 seconds and slow cooled to room temperature (1-2 hours). Cas9 (10 mg/mL, generously gifted by the Seydoux lab) was diluted to 8 μM in storage buffer (20 mM HEPES pH 7.5, 500 mM KCl). Cas9:tr_L_ RNPs were pre-formed by mixing 10 μL 8 uM Cas9 with 10 μL 8 μM tr_L_ and incubating at RT for 10 minutes. RNPs were diluted to 500 pM −1 μM in a 1:1 mix of RNA hybridization buffer and Cas9 storage buffer, then brought to volume with binding buffer and 50 pM substrate in a 20 μL reaction. The binding reaction was incubated at 37° C for 1 hour, then quenched with 2 μL 6X purple loading dye (NEB) at RT. 15 μL of each reaction was immediately loaded onto an 8% TBE gel (Acrylamide/Bis solution 37.5:1, Bio-rad) supplemented with 2 mM MgCl_2_, and run at 4°C at 15W in 1X TBE buffer + 5 mM MgCl2 for 75-90 minutes. The gel was then removed from its casing, wrapped in plastic wrap and exposed to a phosphor screen overnight. The phosphor screen was scanned and imaged using the Typhoon FLA9500 (GE Healthcare).

### *In vitro* cleavage assay

20 μL nuclease-free water, 3 μL NEBuffer 3.1, 3 μL of *in vitro* transcribed 300 nM sgRNA or tr_L_ (produced as described in *in vitro* transcription protocol above) and 1 μL of 1 μM Cas9 nuclease (NEB, M0386S) were mixed and incubated for 10 minutes at 25°C to allow for RNP formation. Next, 3 μL of a 30 nM dsDNA substrate (a 1kb pCN57 amplicon, amplified from pCN57 with oJW134 and oJW1869) was added, mixed thoroughly, pulsed in a microcentrifuge, and incubated at 37°C for 15 minutes. 1 μL of proteinase K (20 mg/mL, Qiagen) was added and samples were incubated at RT for 10 minutes. 6X loading dye was added, and the reactions were run on a 1.5% agarose gel.

### Competition assay

To investigate the fitness costs of ∆tr_L_, cells harboring a naïve wild-type or ∆tr_L_ CRISPR system were co-cultured in a long-term competition assay. Wild-type and ∆tr_L_ cells were inoculated in BHI in triplicate and grown overnight at 37°C, diluted to log phase OD = 0.1, and replicate pairs were mixed 1:1. 50 μL of a 1:100 dilution of this mixture was immediately plated, and 900 μL was frozen at −80°C in 10% DMSO. The remainder of the mixture was grown to stationary phase in the evening, then diluted back 1:1000 and grown overnight. This continued for a total of 5 days, with frozen stocks taken of the cultures each morning before dilution. To determine the ratio of wild-type:∆tr_L_ cells over time, 5 μL of frozen cultures were diluted 1:5000 and plated, and colonies were picked for colony lysis and PCR of the tracrRNA locus (oJW2012/oJW1447). Amplicon length (314 bp for wild-type and 262 bp for ∆tr_L_) allowed us to estimate the relative abundance of each strain in the mixed culture over time.

### Evolutionary analysis

Bacterial strains harboring type II-A CRISPR-Cas systems with previously annotated short-form tracrRNAs^8^ were investigated for evidence of tr_L_ repression. Accession numbers for species investigated are listed in Table S2 along with all metadata compiled in our analysis. The genomic locus between the annotated 3’ end of tracrRNA and the translational start site of Cas9 was interrogated using the DSK k-mer counting software^9^ with k-mer length set to 11 bp and a minimal match frequency of 2. We classified strains as “tr ^+^” if one k-mer instance was on the same strand as tr followed immediately by a 5’-GTTTTA-3’ sequence allowing for 1 mismatch (representing the targeting site within tr_L_) and the other instance was upstream of site 1, immediately followed by a 5’-NGG-3’ sequence allowing for 1 mismatch (representing the targeted site within P_cas9_). We manually inspected the 16 tr_L_^−^ loci with intergenic lengths above 167 bp for candidate tr_L_ targeting sites with a single seed mismatch but perfect lower stems and PAMs, and found one additional trL sequence with a 10 bp match.

We aligned tracrRNA sequences using MAFFT^10^ (default parameters). Cas9 amino acid sequences for these strains were aligned using MUSCLE^11^, and a phylogenetic tree was constructed using FastTree, with the WAG evolutionary model^12^. All multiple sequence alignments and tree generation were performed in Geneious Prime 2020.1.1.

